# Neuronal Transcriptome Disruption, Tau Accumulation and Synapse Loss in Alzheimer’s Knock-in Mice Require Cellular Prion Protein

**DOI:** 10.1101/2023.02.15.528700

**Authors:** Austin Stoner, Li Fu, LaShae Nicholson, Chao Zheng, Takuya Toyonaga, Joshua Spurrier, Will Laird, Zhengxin Cai, Stephen M. Strittmatter

## Abstract

**Background:** Cellular prion protein (PrP^C^) is a high-affinity cell-surface receptor for Amyloid-β oligomers (Aßo). In certain overexpression models of Alzheimer’s Disease (AD), pharmacology and genetics demonstrate its essential role for synaptic plasticity impairment, memory deficits and synapse loss. However, PrP^C^’s role in AD-related phenotypes with endogenous expression levels, its role in tau accumulation and its effect on imaging biomarkers are unknown. The necessity of PrP^C^ for transcriptomic alterations driven by Aß across cell types is unexplored.

**Methods:** The role of PrP^C^ was examined as a function of age in homozygous *App^NL-G-F^/hMapt* double knock-in mice (DKI). Phenotypes of *App^NL-G-F^/hMapt* mice with a deletion of *Prnp* expression (DKI; *Prnp^-/-^*) were compared with DKI mice with intact *Prnp*, mice with a targeted deletion of *Prnp (Prnp^-/-^*), and mice with intact *Prnp* (WT). Phenotypes examined included behavioral deficits, synapse loss by PET imaging, synapse loss by immunohistology, tau pathology, gliosis, inflammatory markers, and snRNA-seq transcriptomic profiling.

**Results:** By 9 months age, DKI mice showed learning and memory impairment, but DKI; *Prnp^-/-^* and *Prnp^-/-^* groups were indistinguishable from WT. Synapse loss in DKI brain, measured by [18F]SynVesT-1 SV2A PET or anti-SV2A immunohistology, was prevented by *Prnp* deletion. Accumulation of Tau phosphorylated at aa 217 and 202/205, C1q tagging of synapses, and dystrophic neurites were all increased in DKI mice but each decreased to WT levels with *Prnp* deletion. In contrast, astrogliosis, microgliosis and Aß levels were unchanged between DKI and DKI; *Prnp^-/-^* groups. Single-nuclei transcriptomics revealed differential expression in neurons and glia of DKI mice relative to WT. For DKI; *Prnp^-/-^* mice, the majority of neuronal genes differentially expressed in DKI mice were no longer significantly altered relative to WT, but most glial DKI-dependent gene expression changes persisted. The DKI-dependent neuronal genes corrected by *Prnp* deletion associated bioinformatically with synaptic function. Additional genes were uniquely altered only in the *Prnp^-/-^*or the DKI; *Prnp^-/-^* groups.

**Conclusions:** A functional *Prnp* gene is required in *App^NL-G-F^/hMapt* double knock-in mice for synapse loss, phospho-tau accumulation and neuronal gene expression. These data support the efficacy of targeting the Aßo-PrP^C^ interaction to prevent Aßo-neurotoxicity and pathologic tau accumulation in AD.

## BACKGROUND

The most common cause of dementia is Alzheimer’s Disease (AD), characterized by the accumulation of amyloid-β (Aβ) plaques and neurofibrillary tangles in the brain [1, 2]. Accumulation of misfolded protein is coupled with synapse loss, inflammation, cognitive decline, and eventually neuronal degeneration and death. Currently, there is no cure for AD, and treatments focus on reducing symptoms and slowing the progression of the disease. Recent clinical trials document some slowing of disease progression during treatment with certain anti-Aß antibodies [3, 4]. Diffusible oligomeric and protofibrillary species of Aß (Aßo) are recognized as being most critical to trigger changes in brain function.

Cellular Prion Protein (PrP^C^) was identified in an unbiased expression cloning experiment as a high-affinity oligomer-specific Aß binding site at the neuronal surface [5–8]. Aßo species interacting with PrP^C^ are detected in mouse models and in human AD brain [9–11]. Numerous studies have examined the requirement for PrP^C^ in mediating effects of Aßo on neurons and mice (reviewed in [8]). Most studies have utilized transgenic mice overexpressing AD-causing variants of APP and PSEN1, and found necessity of PrP^C^ for synapse loss, memory deficits and selective neuronal degeneration using anti-PrP^C^ antibody blockade or PrP antagonists or gene knockout [12–17]. The action of Aßo/PrP^C^ complexes is mediated by aberrant activation of mGluR5 [10, 18–21].

There are limitations and clarifications regarding an Aßo/PrP^C^/mGluR5 model of synaptic damage in AD [8, 22]. First, the *in vivo* data rely largely on transgenic overexpression of Aß and not all transgenic models of AD show PrP-dependence [23]. Whether Aß pathology driven by endogenous expression levels requires this pathway has not been assessed. Second, the transcriptome-wide effect of PrP^C^ on altered gene expression in neurons and glia as a function of AD pathology has not been explored. Third, it is clear that innate immunity and glial reaction have a prominent role in AD [1, 2], but PrP-dependence of interactions between neuronal synaptic mechanisms and glial mechanisms is undefined. Fourth, there is limited or no Tau pathology in the Aß models, so the connection of PrP^C^ and mGluR5 to tauopathy is unclear, though data from iPSC models as well as studies of Fyn and Pyk2 kinases suggest a link. Finally, the role of PrP^C^ in AD-related synaptic loss has not been monitored using clinically relevant biomarkers.

Here, we utilize a homozygous double knock-in (DKI) mouse model with expression from the endogenous *App* and *Mapt* loci (*App^NL-G-F/NL-G-F^, Mapt^hMAPT/hMAPT^* [19, 24, 25] to investigate the effects of constitutive *Prnp* loss on AD-related phenotypes. The DKI strain exhibits memory deficits in the presence of PrP^C^, but not with the *Prnp* null allele. Synapse loss measured in DKI mice by PET or by immunohistology is rescued to wild-type levels by *Prnp* knockout. Similarly, tau accumulation, dystrophic neurites and synaptic tagging by C1q were significantly reduced in DKI mice when PrP^C^ was absent. Transcriptional changes associated with the DKI model were found in multiple cell types as a function of age. *Prnp* deletion normalized DKI-dependent dysregulation of most neuronal genes but few glia DKI-dependent dysregulated genes. Gliosis and Aß plaque in the DKI model were not altered by the absence of *Prnp*. Thus, PrP^C^ is essential for synaptic and neuronal phenotypes in this knock-in AD model, but Aß accumulation and glial reaction are PrP-independent and dissociable from neuronal changes. The findings support therapeutic targeting of the PrP^C^/mGluR5 complex as a neuron-specific approach to ameliorating AD progression.

## METHODS

### Animals

Mice were cared for by the Yale Animal Resource Center and all experiments, including animal husbandry, genotyping, behavioral testing, and euthanasia, were approved by Yale’s Institutional Animal Care and Use Committee (IACUC). Animals were housed in groups of 1–5 mice per cage with access to food and water ad libitum. 12-hour light/dark cycles were maintained throughout the duration of animal housing. All strains were all maintained on a pure C57Bl6J background after more than 10 backcrosses. The *App^NL-G-^ F/NL-G-F, Mapt^hMAPT/hMAPT^* mice were provided by Drs. Saito and Saido, RIKEN Center for Brain Science [19, 24, 25]. The *Prnp^-/-^* mice, Edinburgh strain, were obtained from Dr. Chesebro of the Rocky Mountain Laboratories [12, 26]. We utilized the following abbreviated strain designations:

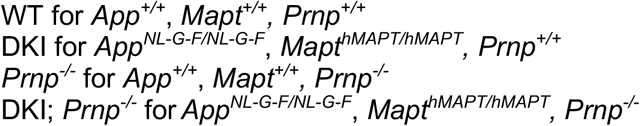

DKI; *Prnp^-/-^* mice were generated by first crossbreeding DKI mice with *Prnp^-/-^* mice to generate triple-heterozygous *App^NL-G-F/+^, Mapt^hMAPT/+^, Prnp^+/-^* mice. The triple-heterozygous mice and their offspring were interbred to create a colony of triple-homozygous DKI; *Prnp^-/-^* mice. Mice were genotyped by Transnetyx (Cordova, Tennessee, USA).

### PET Imaging of Synaptic Density

[^18^F]SynVesT-1 was synthesized at the Yale PET Center as utilized for WT and DKI mouse imaging as described [19, 27]. PET measurements were collected on a Focus 220 (Siemens Medical Solutions). PET data were acquired over the period of 30-60 min post intramuscular injection of [^18^F]SynVesT-1, followed by a transmission scan using ^57^Co. Anesthesia was with isoflurane.

The cerebellum was the reference region for *BP*_ND_ values and calculating SUVR_CB_. Differences in SUVR_CB_ between groups were assessed on a voxel-by-voxel basis in Matlab R2018 (Mathworks) with Statistical Parametric Mapping (SPM12). The Z-score for the difference of two groups was calculated in the framework of the general linear model per voxel. Resulting Z-score maps were projected onto the T2-weighted Ma-Benviste-Mirrione mouse brain template.

### Novel Object Recognition

Mice were tested for novel object recognition as described [13, 20, 21, 28, 29]. Briefly, mice were handled for 5 minutes a day for 3 days prior to testing to familiarize with the experiment and reduce anxiety. Mice were placed in clean, empty, rectangular, covered rat cages to habituate for 1 hour. During acquisition, mice were briefly removed, and 2 identical objects were placed 1 inch from either edge along the long axis of the cage. The objects included: (a wrapped 5mL plastic syringe or single 15mL conical tube with an orange cap (3 months), and a large, black binder clip or large glue stick (9 months)). Familiar object choice was pseudorandom. Mice were placed facing perpendicular to the long axis of the cage and a 10-minute timer was started. Mice were allowed to explore the two identical objects for the duration of the 10 minutes, and the time to accrue 30 total seconds of orofacial object exploration was recorded. The objects were removed and either discarded (15mL conical tube or 5mL wrapped syringe) or wiped down with 70% ethanol. Mice were left in their cages for 1 hour while other mice underwent acquisition.

During testing, one each of the novel and familiar objects were placed on pseudorandom sides of the cage. Orofacial object exploration was timed until 30 total seconds was reached. After testing trials, rat cages were cleaned to eliminate scent cues. The experimenter was blind to novel object identity and genotype. Mice that did not explore both objects or failed to accrue 30 total seconds of orofacial object exploration in less than 6 minutes during either acquisition or testing were removed from the analysis.

### Morris Water Maze

Spatial learning and memory were analyzed using the Morris water maze as previously described [20]. Briefly, testing was performed in a circular, open pool ~1 m in diameter with the water kept at room temperature (~23°C). Throughout testing and analyses, the experimenter was blinded to genotype. Animals were randomly divided into two cohorts of approximately 40 subjects and swims were conducted in 2 sets of 4 days for 7 consecutive days (day 4 of the first cohort overlapping with day 1 of the second cohort). Acquisition trials consisted of 8 attempts per day for 3 consecutive days, divided into one morning and one afternoon training session of four swims each. For each trial, mice were placed in one of four drop zones facing away from the pool’s center in the quadrant opposite the target quadrant, which contained a submerged plexiglass platform. The order of the drop locations varied across each session. The location of the target quadrant was varied in data acquired at 3 months and 9 months of age. Attempts were considered complete once a mouse located and spent 1 second on the platform. On the first day only, mice were allowed to spend 15 seconds on the platform until removal from the pool. Additionally, on the first day only, if a mouse did not find the platform within 60 seconds, it was guided to the platform and allowed to spend 15 seconds on the platform until removal from the pool.

On day 4, the platform was removed for the probe trial assessment. Mice were placed facing away from the center of the pool between the drop zones furthest from the target quadrant and tracked over 60 seconds of swim time. Latency to platform during the training swims and percent time spent in the target quadrant during probe trials were recorded on a JVC Everio G-series camcorder and tracked by Panlab’s Smart software. After completion of the probe trial, a visible platform was placed in the target quadrant and trials were administered until latencies stabilized over a maximum number of 15 trials. The latencies for the final 3 trials were averaged.

### Immunohistology of mouse brain sections

Mice were euthanized by CO2 inhalation and perfused with ice-cold PBS before decapitation and rapid brain dissection. Right brain hemispheres were drop-fixed in 4% paraformaldehyde for 48 hours at 4°C before transfer to PBS at 4°C. Right brain hemispheres were sliced into 40 μm cortical brain sections using a Leica WT1000S vibratome and free-floating sections transferred to PBS with .05% sodium azide for longterm storage.

For AT8 staining, slices were stained using Alexa Fluor 594 Tyramide SuperBoost kit, Streptavidin (Thermofisher; B40935) according to manufacturer instructions. An antigen retrieval step was performed for pThr217 and PSD-95 single staining prior to the blocking step by incubating slices in 1x Reveal Decloaker buffer (Biocare Medical; RV 1000 M) for 15 min at 90°C in an incubator and then 15 minutes at room temperature. For pThr217 staining, sections were incubated in blocking buffer (1% BSA + 1% Triton X-100 in PBS) for 1 h and then incubated in primary antibodies overnight (~24 h) at 4°C in blocking buffer. For all other stains, sections were blocked in 10% normal horse serum (Jackson ImmunoResearch Laboratories) in PBS + 0.2% Triton X-100 for 1 h and then incubated with primary antibodies overnight (~24 h) in 1% normal horse serum in PBS at 4°C.

The following primary antibodies were used: anti-AT8 (Phospho-TAU (Ser202, Thr205), Biotin; Invitrogen MN1020B; 1:100), anti-LAMP-1 (1D4B) (lysosome-associated membrane protein 1; Santa Cruz sc-19992; 1:500) anti-IBA1 (ionized calcium-binding adapter molecule 1; Wako 019-19741; 1:250), anti-PSD-95 (for PSD-95 single stain) (postsynaptic density protein 95; Invitrogen 51-6900; 1:250), anti-PSD-95 (for PSD- 95/C1Q co-stain) (Millipore MAB1596; 1:200), anti-SV2A (synaptic vesicle glycoprotein 2A; Abcam 32942; 1:250), anti-NEUN (Fox-3; Millipore ABN91; 1:500), anti-C1Q (Abcam ab182451; 1:1000), anti-pThr217 (phospho-TAU threonine 217; Invitrogen 44-744; 1:200), anti-pS396 (phospho-TAU serine 396; Invitrogen 44-752G; 1:250), anti-PrP (8H4) (prion protein; Abcam ab61409; 1:1000), anti-GFAP (glial fibrillary acidic protein; Abcam ab4674; 1:2000).

Sections were washed in PBS three times for 5 min each and incubated in secondary antibodies in PBS (goat or donkey anti-rabbit, anti-mouse, or anti-rat fluorescent antibodies; Invitrogen Alexa Fluor; 1:500) for 1 h at room temperature in the dark. Sections were washed in PBS three times for 5 min each and then mounted onto glass slides (Superfrost Plus, Fisher Scientific) and cover-slipped with Vectashield (Vector Laboratories H-1200) antifade aqueous mounting medium.

For Thioflavin S (Sigma; T1892) staining, slices were washed twice for 5 min with 70% ethanol, stained with 0.1% ThioS in 70% ethanol for 15 min at room temperature, and washed twice for 5 min with 70% ethanol before being mounted as previously described. For pThr217 staining, after the three post-secondary PBS washes, slices were briefly dipped in ddH2O and then incubated in CuSO4 buffer (10 mM CuSO 4 in 50mM ammonium acetate buffer, pH 5) for 15 min at room temperature. Slices were then briefly dipped again in ddH2O, returned to PBS, and mounted as previously described.

### Imaging and analysis of immunohistochemistry

Staining for synaptic proteins with anti-SV2A and anti-PSD-95 antibodies was imaged using a Zeiss LSM 800 confocal microscope with a 63X 1.4 NA oil-immersion lens. The percent area occupied by immunoreactive synaptic puncta from the polymorphic layer of the dentate gyrus was measured as described [12, 19]. For imaging of the tissue stained for AT8, NeuN, GFAP, IBA1, CD68, pS396, pThr217, Lamp1, Aß, and Thioflavin S, a Zeiss LSM 800 confocal microscope with a 20X 0.8 NA air-objective lens was used. Imaged regions include hippocampus, medial cortex, or lateral cortex. pS396 was imaged in medial and lateral cortex. GFAP, Iba1, CD68, Thioflavin S, and Aß were imaged using a 5×3 tile and *z-* stack through the full slice in the center of the hippocampus with maximum intensity projection. pThr217 was imaged using a 2×2 tile and *z*-stack through the full slice in CA1 with maximum intensity projection. AT8 and Lamp1 were imaged using a 2×2 tile and *z*-stack through the full slice in medial cortex with maximum intensity projection. The percent area occupied by immunoreactive signal was quantified using ImageJ software. Statistical analysis was based on separate mice, with two slices averaged per mouse [19].

C1q synaptic targeting and PrP/C1q colocalization experiments were completed as described [19]. Briefly, images were acquired using a Zeiss LSM 800 confocal microscope with Airyscan detector and a 63X 1.4 NA oil-immersion lens. 5-layer z-stacks were acquired from three adjacent locations in CA1 per slice. Images were acquired using a magnification factor of 3x and optimized XY pixel size and Z pixel dimension per Zeiss Zen software (~34 nm and 200 nm, respectively). Following image capture, images were 3D Airyscan processed in Zeiss Zen Blue with an Airyscan filter strength of 6. C1q synaptic targeting and PrP/C1q colocalization were calculated using the JACOP plug-in in ImageJ [30].

Immunofluorescent staining of brainstem tissue utilized tissue from 20-month-old mice. After 24 hrs of post-fixation, tissue was cut through the midline and sagittally sectioned into 50 μm free-floating sections by vibratome. Sections were collected and stored in serial order. Sections were blocked with PBS containing 0.1% Triton X-100 (American Bio; AB02025) and 1% BSA for 1h at room temperature. Sections were incubated in rabbit anti-Tyrosine Hydroxylase (Millipore AB152; 1:500) primary antibody diluted in blocking buffer overnight at 4°C. Slices were washed with PBS and then incubated with goat anti-rabbit Alexa Fluor488 (Thermo Fisher Scientific #A-11008, 1:500) secondary antibody for 1 h at room temperature in the dark. After washing in PBS, sections were dipped into distilled water and then incubated in ammonium acetate with 10 mM CuSO4 (pH 5) for 15 min at room temperature to minimize lipofuscin autofluorescence. Samples were then washed in PBS and incubated with DAPI (1:1000) for 1 hour at room temperature. After further washing in PBS, sections were mounted onto glass slides (Superfrost Plus, Fisher Scientific) and coverslipped with Prolong Diamond (ThermoFisher Scientific; P36965) antifade aqueous mounting medium. Tyrosine hydroxylase sections were imaged with a Leica DMi8 confocal microscope with a 10X 0.4 numerical aperture air-objective lens. Seven μm *z*-stacks were acquired from locus coeruleus. Six image slices were *z*-stack projected via maximum orthogonal projection.

All images were acquired, processed, and analyzed by an experimenter blinded to animal genotypes. Target marker’s fluorescence intensity, and immunoreactive area were quantified using Image J with macros. Positive TH cell numbers were counted blindly to animal genotypes.

### Immunoblotting

Mice at 20-months’ age were sacrificed via rapid decapitation and hemispheres separated medially on ice using a sharp razor blade. Cortex were microdissected and homogenized on ice with polypropylene pellet pestles in three times the brain tissue weight in RIPA lysis buffer (Millipore 20-188) containing PhosSTOP (Roche; 4906845001) and cOmplete-mini protease inhibitor cocktail (Roche; 11836153001) to extract cytosolic proteins. After centrifugation for 45 minutes at 100,00 × *g* at 4°C, the supernatants were collected and boiled in Laemmli sample buffer with 5% β-mercaptoethanol and 1% SDS at 95°C for 5min. Sample volumes were normalized to total protein concentration by BCA assay (Thermo Scientific; 23225).

Supernatants were then subjected to SDS-PAGE through 4–20% Tris-glycine gels (Bio-Rad; 5671095) and transferred onto nitrocellulose membranes (Invitrogen #IB23001) using an iBlot 2 Gel Transfer Device (Invitrogen; IB21001). Actin was run on the same gel as the loading control. Nitrocellulose membranes were blocked in blocking buffer for fluorescent western blotting (Rockland; MB-070-010) for 1 hour at room temperature and incubated overnight in primary antibodies at 4°C. Anti-pTau S202/ T205 (AT8) (Thermo Fisher Scientific #MN1020, 1:700), anti-C1Q (Dako A0136; 1:1000), anti-beta Actin (D6A8) Rabbit mAb (Cell Singnaling 8457S; 1:2000) and anti-beta Actin (Abcam ab8226; 1:5000) were used as primaries in this experiment.

Membranes were washed 3 times 5 mins each with TBST after primary antibody incubation and then incubated with appropriate secondary antibodies for 1h at room temperature. Anti-Mouse IRDye 800CW (LI-COR Biosciences #926–32,212, 1:8000) and α-rabbit IRDye 680CW (LI-COR Biosciences; 926–68,023, 1:8000) were used as secondary antibodies. Immunoblots were visualized with a LI-COR Odyssey infrared imaging system. Quantification of band intensities was performed within a linear range of exposure and by ImageJ software. Protein levels were normalized to beta-actin level.

### Single nuclei 10x genomic sequencing and analysis

Mouse brain tissue collection and mouse brain nuclei isolation for single nuclei RNA-seq were completed as previously described [19]. Briefly, WT, *Prnp^-/-^*, DKI, and DKI; *Prnp^-/-^* mice were aged to 40.3±0.7 weeks (10 months) or 143.9±3.5 weeks (20 months) then sacrificed via rapid dissection. Cortical and hippocampal regions from the left-brain hemisphere were microdissected, pooled and immediately frozen on dry ice then stored at −80°C until nuclei isolation. Sample tissue (50-100 mg wet weight) was homogenized, placed on top of homogenization buffer, and centrifuged for 1h. Nuclei pellets were obtained, resuspended, and counted on a hemocytometer to inform subsequent dilution or concentration to 700-1200 nuclei/μl for generating single nuclei cDNA libraries.

Barcode incorporated single-nucleus cDNA libraries were constructed using the Chromium Single Cell 3’ Reagents Kit v3 (10x Genomics) following the manufacturer’s guidelines. Sample Libraries, for both 10- and 20-month-aged cohorts were then pooled and underwent batch sequencing on an Illumina NovaSeq 5000 using single indexed paired-end HiSeq sequencing. A sequencing depth of >300 million reads was achieved for all samples with an average read depth of 30,000 reads per nuclei. The resulting sequencer BCL files were demultiplexed into FASTQ files. Sequenced samples were then aligned to the mm10-2020-A *Mus musculus* reference genome using the Cell Ranger Count software (pipeline version 6.01, 10x Genomics), generating barcoded sparse matrices of gene-nuclei raw UMI counts.

Sample gene count matrices were converted and combined into a single anndata object for quality control (QC) and downstream processing with Scanpy (version 1.8.2) [31], a Python-based gene expression analysis toolkit. Genes detected in less than 10 nuclei were discarded. Nuclei with over 5% of UMI counts mapped to mitochondria genes as well as nuclei with less than 50 genes or more than 8000 genes detected were considered outliers and also discarded. In order to perform comparative gene expression analysis, the retained UMI counts were normalized to their library size by scaling the total number of transcripts to 10,000 per nuclei then Log-transformed. In total, across all experimental groups, 206,575 (10 months, n=17 samples) and 219,900 (20 months, n=23 samples) nuclei and 27,479 genes were retained post QC processing and clustering.

Sample snRNA-seq data were integrated for clustering using Scanpy’s implementation of *Seurat_v3* [32]. This process first finds nuclei expression state ‘integration anchors’ across sample datasets by identifying common highly variable genes (HVGs) using the *highly_variable_genes* function of Scanpy. In brief, using raw UMI counts, HVGs were identified by ranking the computed normalized variance of each gene across all nuclei. The set of HVGs identified within each sample dataset were then merged to remove batch-specific genes, after which the top 2000 HGVs were annotated within the anndata object. Next, the regression of total library size and mitochondrial transcripts per nuclei was performed with sample-based batch correction using the *Combat* function in Scanpy. The *Scale* function was used to scale the corrected expression matrix to a max unit variance of 10 standard deviations. A neighborhoods graph of the corrected matrix was constructed using the *Neighborhoods* function, with a knn restriction of 10 and the first 25 principal components, computed from HVGs, as input parameters. Integrated Leiden clustering of nuclei into cell-type subgroups was performed from the neighborhoods graph and dimensionally reduced in UMAP space for visualization. To refine the clustering, nuclei doublets were removed from the dataset and integrated Leiden clustering was reiterated. Nuclei were deemed doublets if associated with clusters enriched for cell-specific marker genes of multiple cell types within the same cluster, indicating mixed-cell expression profiles.

Nuclei clusters enriched with a particular set of marker genes were considered to be of the corresponding primary cell type. Using the *rank_genes_groups* function, the top highly differentially expressed marker genes for individual Leiden clusters were tested against the rest using the Wilcoxon Ranked-Sum test. The highest-ranked genes with singular cluster enrichment were deemed marker genes and their cell specificity was varied by literature to determine cluster cell types.

The *rank_genes_groups* function, with Wilcoxon Ranked-Sum test, was also used for differential expressed gene (DEG) analysis between experimental groups for each identified cell type. Only genes expressed in a minimum of 5% of all nuclei post-ranking were considered differentially expressed. Significant DEGs were defined as genes with an adjusted *p-value* (false discovery rate) less than 0.005 and Log-transformed fold change greater than ±0.25 (Supplemental Table S1).

Gene set enrichment analysis of DEGs from specific cell types was performed using the network visualization software, Cytoscape (version 3.9.1) [33], with application extensions ClueGo (version v2.5.9) [34] and STRING (version 2.0.0) [35]. DEG lists were queried in ClueGo for the overrepresentation of functional and pathway term associations in Gene Ontology (GO) Molecular Functions, Reactome and KEGG ontology databases. A two-sided hypergeometric test with BH-adjusted p-value of 0.0005 and 4-gene threshold was used for term enrichment selection. GO fusion of represented terms was used to collapse redundant and related biological themes with more than 50% similarity of associated genes. Filtered term associations were organized as degree-sorted networks for optimal visualization. The same gene sets were separately tested for cellular localization enrichment using GO cellular compartments and protein-protein interaction (PPI) networks using STRING. Highly enriched PPI networks were selected using a confidence score of 0.4 for strength of gene associations with MCL clustering.

### Experimental design and statistical analysis

One-way ANOVA with post hoc Tukey’s multiple comparisons tests, One-way ANOVA with post hoc Dunnett’s multiple comparisons tests, and unpaired two-tailed *t*-tests were performed using GraphPad Prism software, version 9. Group means ± SEM and sample sizes (*n*) are reported in each figure legend. Data were considered statistically significant if *p*<0.05. For all figures, all statistically significant group differences are labeled. For any given group comparison, the absence of any indication of significant difference implies lack of significance by the applied statistical test.

## RESULTS

### Aged *DKI* mouse learning and memory deficits require PrP^C^

We sought to evaluate whether *Prnp* deletion would rescue behavioral deficits in the DKI model. DKI mice exhibited no significant spatial learning or memory deficits as measured by Morris water maze at 3 months age (Fig. 1A, B), which is consistent with prior studies [19]. In a separate memory test for object recognition, DKI mice showed a similar preference for the novel object as did WT, *Prnp^-/-^*, DKI and DKI; *Prnp^-/-^* mice at 3 months age (Fig. 1C). Thus, at this young adult age, DKI mice do not demonstrate significant memory deficits as measured by these two tests.

**Figure 1.**
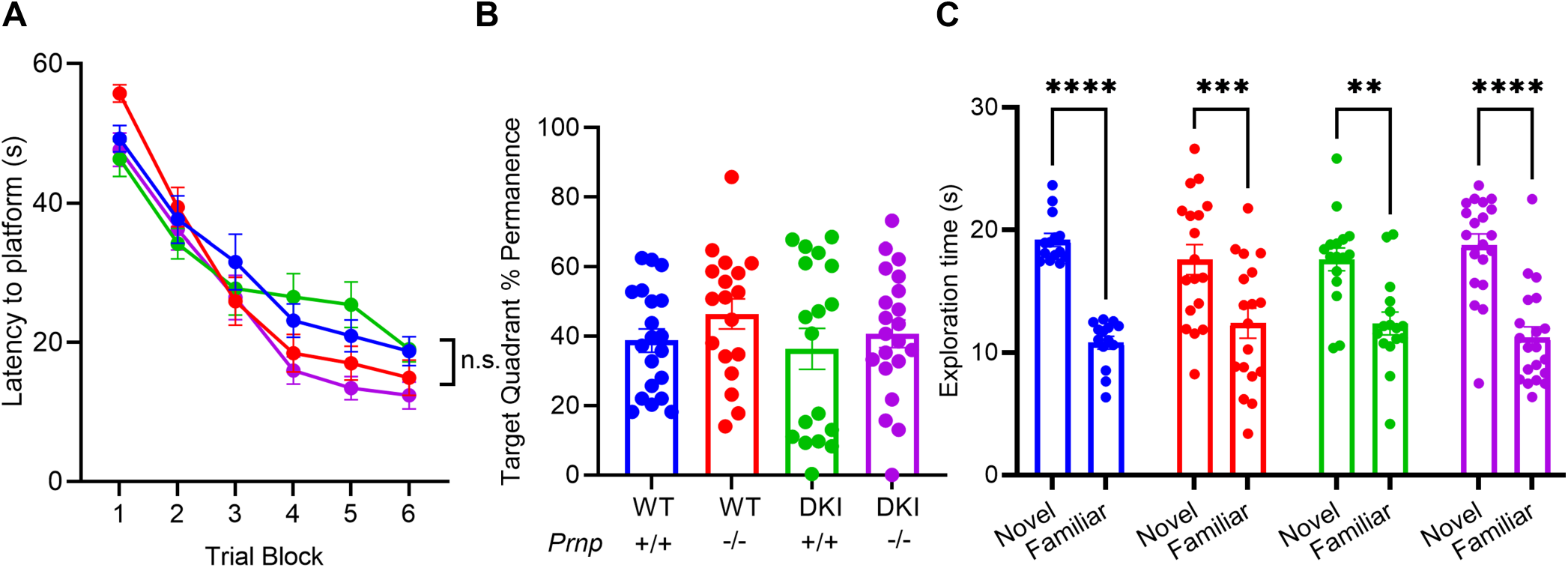
Young Adult DKI mice demonstrate no learning and memory deficit at 3 months. (A) 3-month-old WT, *Prnp^-/-^*, DKI and DKI; *Prnp^-/-^* mice completed the MWM to investigate the age of spatial memory deficit. Latency is defined as the average time of 4 trials to find a hidden platform across 6 acquisition sessions. No significant difference was observed in the time to reach the platform during the final acquisition session in any genotype compared to DKI. Data are graphed as mean ± SEM, analyzed by two-way ANOVA with Dunnett’s multiple comparisons test, P>0.05, n=20 for WT, n=18 for *Prnp^-/-^*, n=18 for DKI, and n=21 for DKI; *Prnp^-/-^*. (B) 24h after completing the final acquisition session, all 4 genotypes completed a probe trial. The percent permanence is defined as the fraction of 60s spent in the quadrant where the platform was during acquisition trials. No significant difference was observed in the percent permanence in any genotype compared to DKI. Data are graphed as mean ± SEM, analyzed by two-way ANOVA with Dunnett’s multiple comparisons test, P>0.05, n=20 for WT, n=18 for *Prnp^-/-^*, n=18 for DKI, and n=21 for DKI; *Prnp^-/-^*. (C) 3-month-old WT, *Prnp^-/-^*, DKI and DKI; *Prnp^-/-^* mice completed the NOR test. All 4 genotypes preferred to interact with the novel object compared to the familiar object. Data are graphed as mean ± SEM, analyzed by two-way ANOVA with Sidak’s multiple comparisons test, ** P<0.01, ***P<0.001, ****P<0.0001, n=14 for WT, n=18 for *Prnp^-/-^*, n=16 for DKI, and n=20 for DKI; *Prnp^-/-^*.

DKI mice have been shown to demonstrate spatial memory deficits as measured by Morris water maze by 12 months age [19]; however, we sought to test if these mice exhibit observable deficits at an earlier age.

We conducted the Morris water maze at 9 months of age on the mice previously tested at 3 months, with the target quadrant placed opposite the 3-month location. DKI mice exhibited learning deficits compared to WT mice, taking significantly longer to find a hidden platform in the final training block (Fig. 2A, *p*<0.05). These mice also demonstrated an impaired ability to recall the target quadrant location in the probe trial relative to WT mice (Fig. 2B, *p*<0.05). Both these observed deficits in DKI mice were rescued by *Prnp* gene knockout. All 4 genotypes demonstrated an equivalent latency to locate a visible platform (Fig. 2A), suggesting the differences observed were attributed to spatial memory deficits rather than vision or motor impairment.

**Figure 2.**
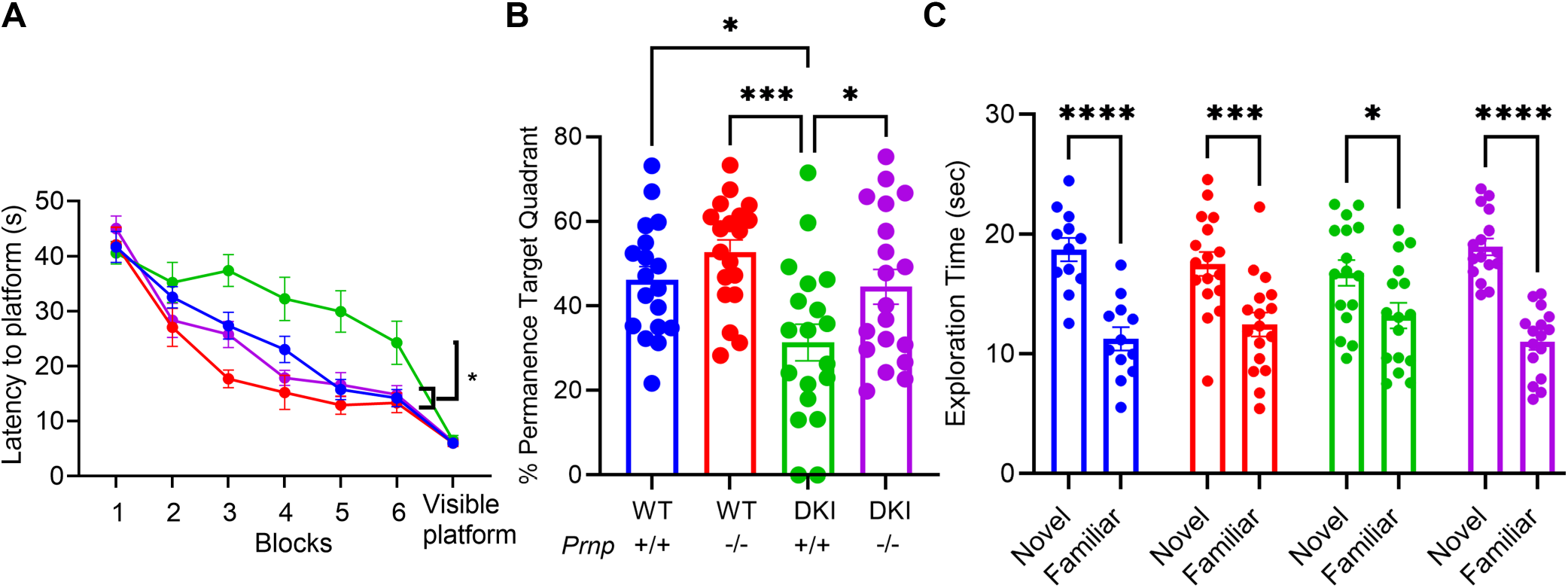
Aged DKI mouse learning and memory impairment at 9 months requires *Prnp* expression. (A) 9-month-old WT, *Prnp^-/-^*, DKI and DKI; *Prnp^-/-^* mice repeated the MWM to assess the onset of spatial memory deficit. Latency is defined as the average time of 4 trials to find a hidden platform across 6 acquisition sessions. DKI mice took significantly longer to reach the platform during the final acquisition session compared to WT, *Prnp^-/-^* and DKI; *Prnp^-/-^*. The DKI; *Prnp^-/-^* mice showed learning equal to WT levels. To rule out the contribution of visual or motor impairments, latency to a visible platform was measured and no significant difference was observed in any genotype compared to WT. Data are graphed as mean ± SEM, analyzed by two-way ANOVA with Dunnett’s multiple comparisons test, *P<0.05, n=18 for WT, n=19 for *Prnp^-/-^*, n=19 for DKI, and n=19 for DKI; *Prnp^-/-^*. (B) 24h after completing the final acquisition session, all 4 genotypes completed a probe trial. The percent permanence is defined as the fraction of 60s spent in the quadrant where the platform was during acquisition trials. DKI mice spent significantly less time in the target quadrant compared to WT, *Prnp^-/-^* and DKI; *Prnp^-/-^* mice. The DKI; *Prnp^-/-^* mice showed rescue equal to WT levels. Data are graphed as mean ± SEM, analyzed by ordinary one-way ANOVA with Dunnett’s multiple comparisons test, *P<0.05, ***P<0.001, n=18 for WT, n=19 for *Prnp^-/-^*, n=19 for DKI, and n=19 for DKI; *Prnp^-/-^*. (C) 9-month-old WT, *Prnp^-/-^*, DKI and DKI; *Prnp^-/-^* mice repeated the NOR test. All 4 genotypes preferred to interact with the novel object compared to the familiar object, though DKI; *Prnp^-/-^* mice exhibited a trend towards less novel object recognition. Data are graphed as mean ± SEM, analyzed by two-way ANOVA with Sidak’s multiple comparisons test, * P<0.05, ***P<0.001, ****P<0.0001, n=12 for WT, n=17 for *Prnp^-/-^*, n=16 for DKI, and n=16 for DKI; *Prnp^-/-^*.

We also assessed novel object recognition in these mice at 9 months of age. DKI mice continued to show a significant preference for the novel object that was not statistically different from the other groups (Fig. 2C); however, we observed a trend toward decreased novel object recognition in this group relative to the other groups.

Thus, we observed learning and memory deficits in 9 months old DKI mice during Morris water maze that were fully rescued by *Prnp* deletion. Novel object recognition was not sufficiently impaired in DKI mice at 9 months to assess a requirement for PrP^C^.

### *Prnp* deletion prevents synapse loss in DKI mice

Synapse loss is an early event in AD progression that correlates strongly with cognitive decline [36, 37]. Thus, we sought to understand if the behavioral deficits observed in DKI mice and subsequent rescue with *Prnp* knockout correspond with changes in synapse density. DKI mice have been shown to exhibit detectable decreases in synapse density at 12 months of age using SV2A PET [19]. At 10 months age, we observed reduced [^18^F]SynVesT-1 binding in the hippocampus of DKI mice relative to WT mice (Fig. 3A,C, *p*<0.05). Meanwhile, DKI; *Prnp^-/-^* mice demonstrated a full rescue in hippocampal [^18^F]SynVesT-1 standardized uptake value ratio SUVR_CB_ back to WT levels (Fig. 3A,C). We also observed that *Prnp^-/-^* without DKI background showed significant deficits in [^18^F]SynVesT-1 binding in the olfactory bulb region compared to WT mice (Fig. 3A). This data supports previous studies that have implicated PrP^C^ in maintaining mature olfactory sensory neurons [38–40]. Interestingly, despite observing this effect on a wildtype background, DKI; *Prnp^-/-^* mice did not demonstrate synaptic density decrease in olfactory bulb compared to WT mice (Fig. 3A).

**Figure 3.**
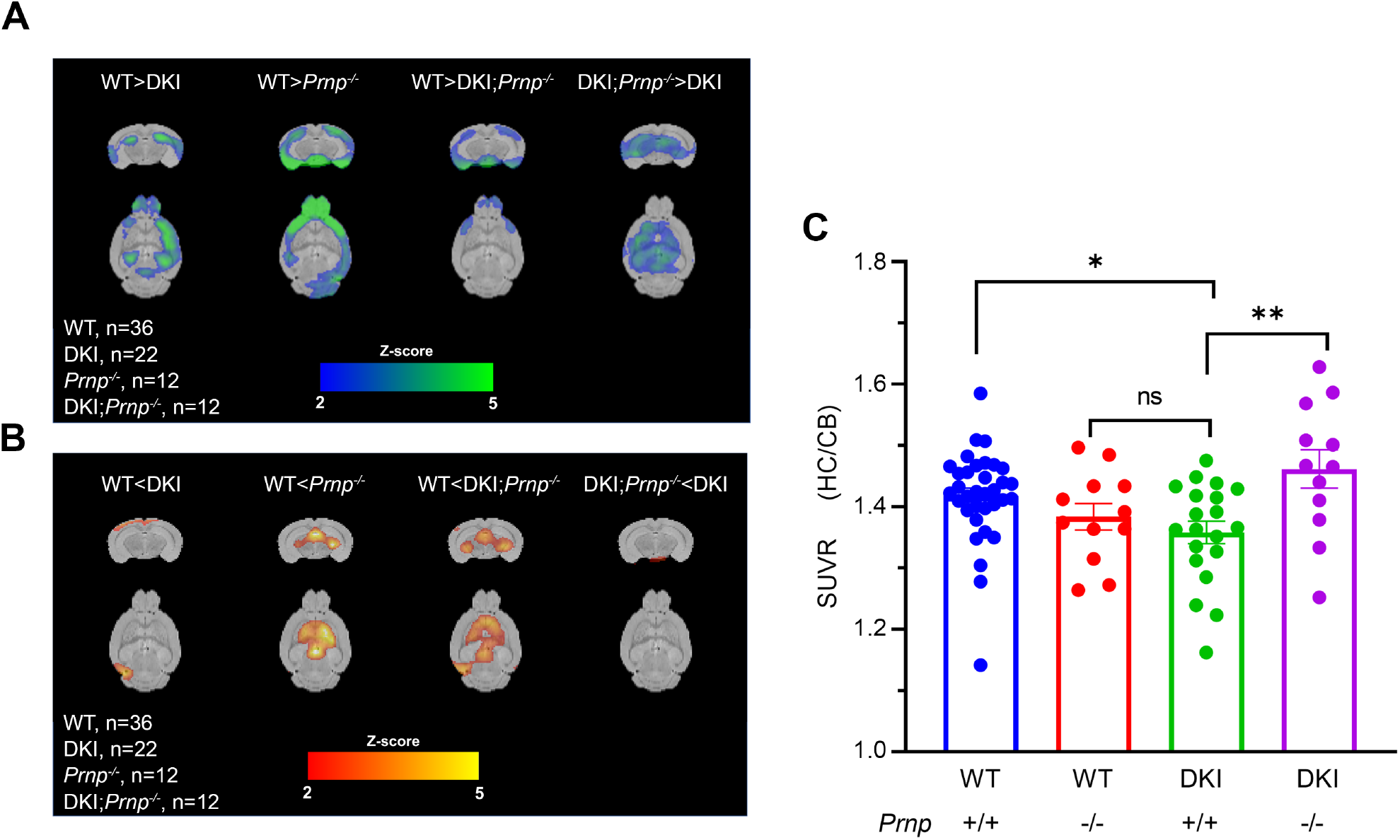
*Prnp* deletion rescues DKI-dependent synapse density reduction measured by SV2A PET. (A-B) Voxel-wise analysis t value map comparing averaged [18F] SynVesT-1 PET in 15-month-old mouse brains. Activity is expressed as SUVR (normalized to cerebellum). Color scale represents magnitude of difference in activity in that genotypic comparison. (C) ROI-based comparison of hippocampal [18F]SynVesT-1 SUVR (normalized to cerebellum) in WT, *Prnp^-/-^*, DKI and DKI; *Prnp^-/-^* mice. DKI mice demonstrate decreased hippocampal synaptic density compared to WT, while *Prnp* knockout restores hippocampal synapse density in the DKI background. Data are graphed as mean ± SEM, analyzed by ordinary one-way ANOVA with Dunnett’s multiple comparisons test, * P<0.05, **P<0.01, n=36 for WT, n=12 for *Prnp^-/-^*, n=22 for DKI, and n=12 for DKI; *Prnp^-/-^*.

We also tested whether *Prnp* loss prevented synapse loss by analyzing synaptic marker immunostaining. 10-month-old DKI mice exhibited significant (*p*<0.05) reduction in the presynaptic marker SV2A in dentate gyrus (DG) compared to WT mice, while DKI; *Prnp^-/-^* mice demonstrated a full-rescue to WT levels (Fig. 4A, B). At 10 months age, DKI mice did not exhibit a statistically significant decrease in the postsynaptic marker postsynaptic density protein 95 (PSD-95) in DG compared to WT, though we did observe a trend towards a DG PSD-95 immunoreactivity increase in DKI; *Prnp^-/-^* mice compared to DKI mice (Fig. 4C,D).

**Figure 4.**
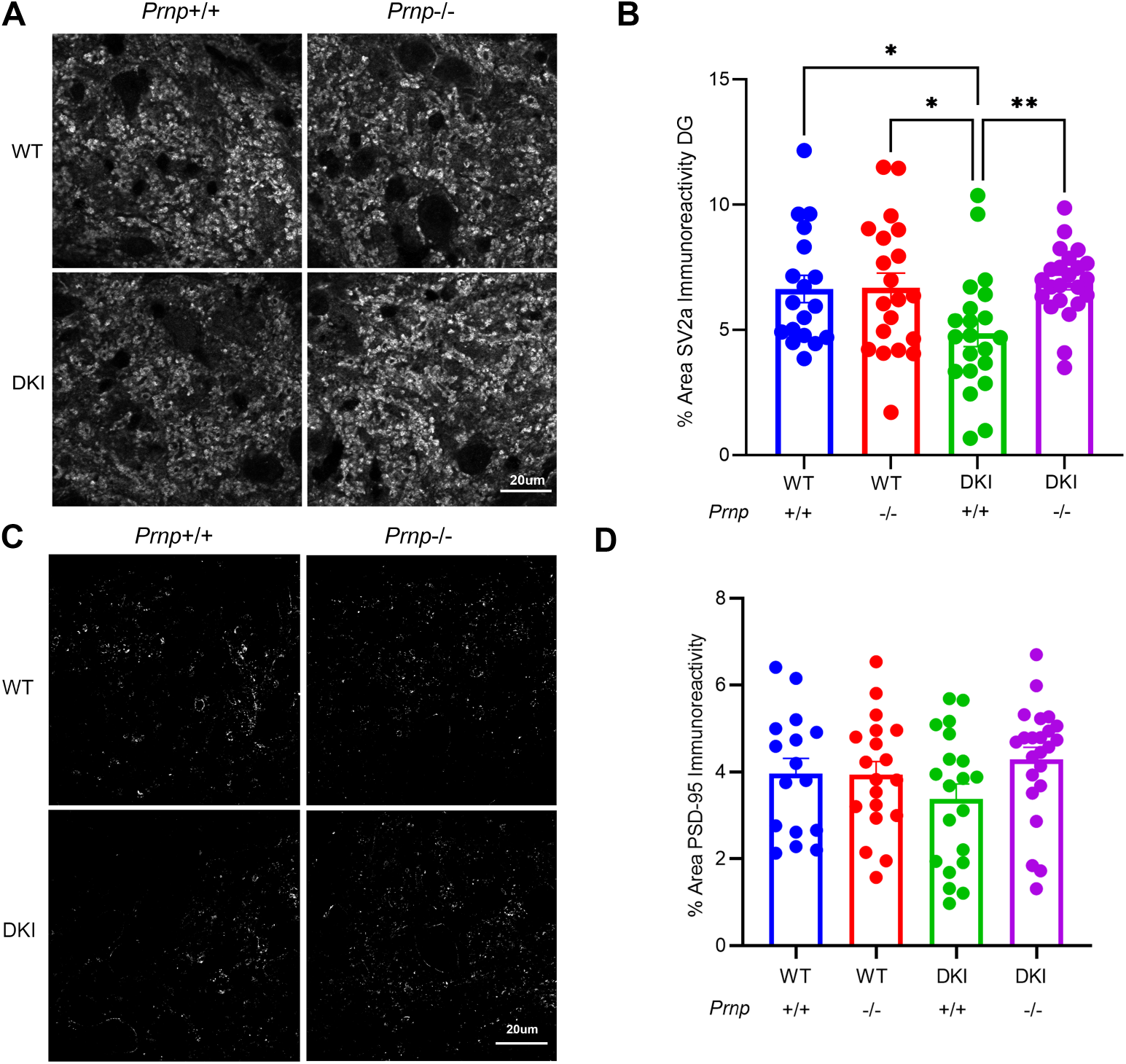
*Prnp* deletion reverses DKI-dependent loss of synaptic immunohistochemical markers. (A) Representative images of staining for the pre-synaptic marker SV2A from the dentate gyrus of 10-month-old WT, *Prnp^-/-^*, DKI and DKI; *Prnp^-/-^* mice. Scale bar = 20 μm. (B) Quantification of SV2A immunoreactive area in the dentate gyrus demonstrates a significant decrease in synapse density in DKI mice that is not observed in DKI; *Prnp^-/-^* mice. Data are graphed as mean ± SEM, analyzed by ordinary one-way ANOVA with Dunnett’s multiple comparisons test to DKI, *P<0.05, **P<0.01, n=18 for WT, n=20 for *Prnp^-/-^*, n=21 for DKI, and n=23 for DKI; *Prnp^-/-^*. (C) Representative images of staining for the post-synaptic marker PSD-95 from the dentate gyrus of 10-month-old old WT, *Prnp^-/-^*, DKI and DKI; *Prnp^-/-^* mice. Scale bar = 20 μm. (D) Quantification of PSD-95 immunoreactive area in the dentate gyrus demonstrates a trend (P= 0.09) towards greater area in DKI; *Prnp^-/-^* mice compared to DKI mice. Data are graphed as mean ± SEM, analyzed by ordinary one-way ANOVA with Dunnett’s multiple comparisons test to DKI, n=16 for WT, n=19 for *Prnp^-/-^*, n=20 for DKI, and n=23 for DKI; *Prnp^-/-^*.

Monoaminergic axon and cell degeneration is a pathologic hallmark of AD and is observed in some mouse models [12, 41, 42]. We assessed brain stem neuronal loss in 20-month-old DKI mice using tyrosine hydroxylase (TH) immunoreactivity. We observed a non-significant trend of decreased TH immunoreactivity but not cell number in the locus coeruleus of DKI mice compared to WT (Suppl. Fig. S1). Since there was no robust degeneration phenotype in DKI mice, a role of *Prnp* could not be assessed.

### Phospho-tau accumulation in DKI mice is reduced by *Prnp* knockout

The accumulation of hyperphosphorylated Tau and neurofibrillary tangles are pathological hallmarks of AD triggered by Aß pathology [1, 2, 43]. The DKI model replaces murine *Mapt* with human *MAPT* and demonstrates increased levels in certain Tau epitopes, including AT8, pThr217, and pS396 [19, 25]. We assessed the influence of *Prnp* expression on phosphorylated Tau levels in DKI mice by immunohistochemistry and biochemistry. In 10-month-old animals, we detected a significant increase in AT8 immunoreactivity in medial cortex (*p*<0.0001) and pThr217 immunoreactivity in the CA1 region of hippocampus (*p*<0.0001) for DKI compared to WT (Fig. 5A-D). The DKI; *Prnp^-/-^* tissue showed a significant decrease of both AT8 and pThr217 immunoreactivity relative to DKI, with pThr217 immunoreactivity indistinguishable from WT levels (Fig. 5A-D). Biochemical analysis of 20-month-old mice cortical tissue revealed significantly elevated AT8 level in DKI mice relative to WT, with a significant decrease of DKI; *Prnp^-/-^* levels relative to DKI to a level equaling WT values (Fig. 5E-F). Thus, the accumulation of these AD-associated phospho-Tau epitopes depends on PrP^C^ in DKI mice.

**Figure 5.**
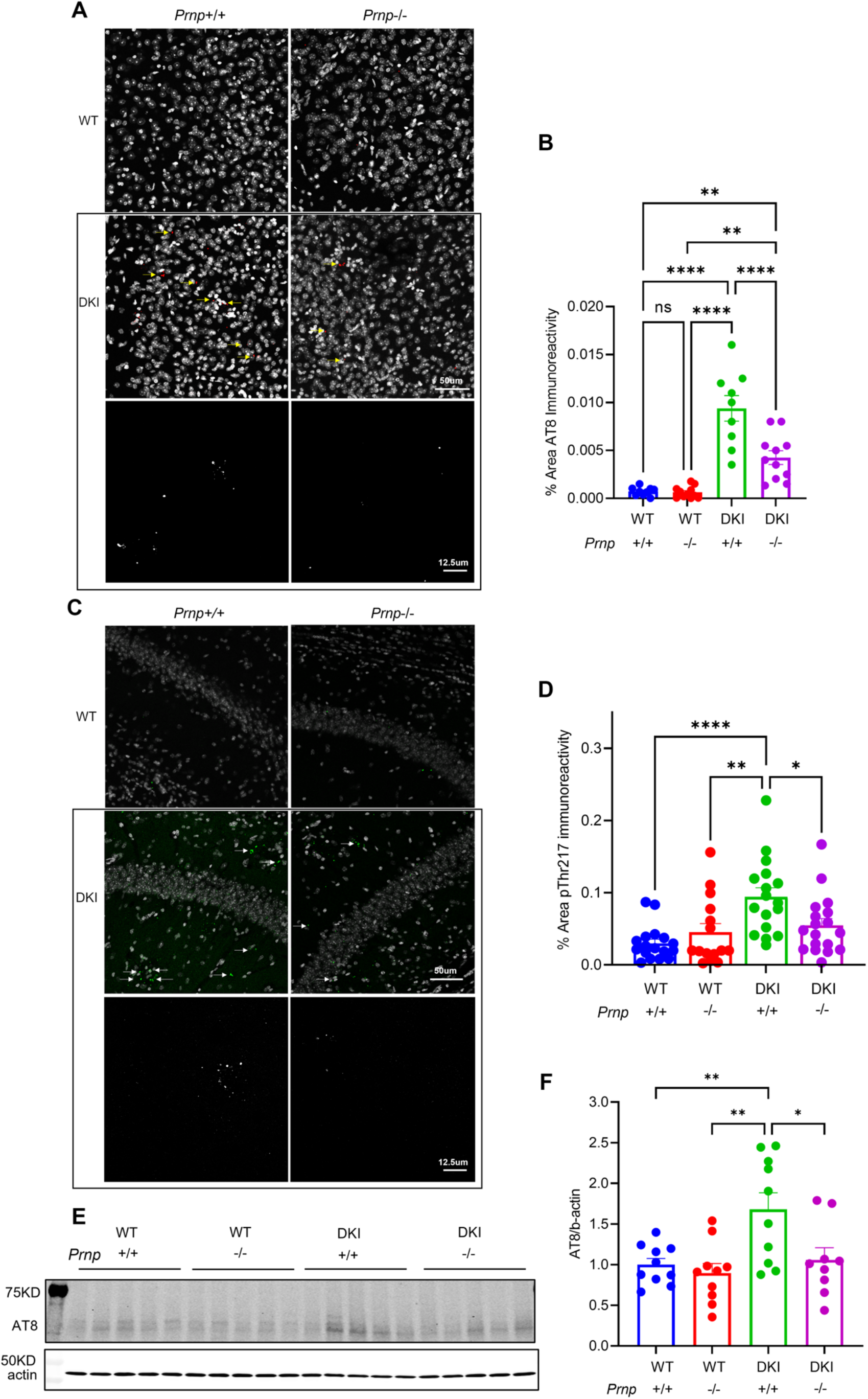
Phospho-tau accumulation in DKI mice is reduced by *Prnp* knockout. (A) Representative images of AT8 (red) staining in the medial cortex of 10-month-old old WT, *Prnp^-/-^*, DKI and DKI; *Prnp^-/-^* mice. Gray is DAPI signal, and AT8 inclusions are indicated by yellow arrows in the upper four panels with scale bar = 50 μm. The lower two panels show the AT8 signal only in gray scale at higher magnification with scale bar = 12.5 μm. (B) Quantification of AT8 immunoreactive area in medial cortex demonstrates a significant increase in phospho-tau accumulation in DKI mice compared to WT. There is a significant decrease in DKI; *Prnp^-/-^* mice relative to DKI. Data are graphed as mean ± SEM, analyzed by ordinary one-way ANOVA with Tukey’s multiple comparisons test, **P<0.01, ****P<0.0001, n=9 for WT, n=11 for *Prnp^-/-^*, n=9 for DKI, and n=11 for DKI; *Prnp^-/-^*. (C) Representative images of pThr217 (green) staining in the CA1 of hippocampus of 10-month-old WT, *Prnp^-/-^*, DKI and DKI; *Prnp^-/-^* mice. Gray is DAPI signal, and pThr217inclusions are indicated by arrows in the upper four panels with scale bar = 50 μm. The lower two panels show the pThr217 signal only in gray scale at higher magnification with scale bar = 12.5 μm. (D) Quantification of pThr217 immunoreactive area in CA1 demonstrates a significant increase in phospho-tau accumulation in DKI mice compared to WT. The DKI; *Prnp^-/-^* samples show a significant decrease relative to DKI. Data are graphed as mean ± SEM, analyzed by ordinary one-way ANOVA with Dunnett’s multiple comparisons test, *P<0.05, **P<0.01,****P<0.0001, n=17 for WT, n=15 for *Prnp^-/-^*, n=17 for DKI, and n=18 for DKI; *Prnp^-/-^*. (E) Immunoblot image of cortical brain lysates stained for AT8 from 20-month-old WT, *Prnp^-/-^*, DKI and DKI; *Prnp^-/-^* mice. (F) Quantification of AT8 antibody immunoreactive protein levels by densitometric analysis. The level of Tau phosphorylated at S202/T205 (AT8) is significantly greater in DKI mice compared to WT or *Prnp^-/-^*. The DKI; *Prnp^-/-^* AT8 level is less than that from DKI. Data are graphed as mean ± SEM, analyzed by ordinary one-way ANOVA with Dunnett’s multiple comparisons test, *P<0.05, **P<0.01, n=10 for WT, n=10 for *Prnp^-/-^*, n=10 for DKI, and n=9 for DKI; *Prnp^-/-^*.

The DKI mice also exhibited increased pS396 immunoreactivity in cerebral cortex (*p*<0.0001) compared to WT (Suppl. Fig. S2A-D) [19]. However, the enhanced pS396 signal in DKI mice co-localizes exclusively with a marker for oligodendrocytes (Olig2) rather than a neuronal marker (NeuN) (Suppl. Fig. S2E, F). The effect of *Prnp* deletion was complicated in these non-neuronal cells. In cerebral cortex, there was a slight increase for *Prnp^-/-^* mice in oligodendrocyte-lineage cells. For the DKI; *Prnp^-/-^* samples, the levels were similar to DKI (Suppl. Fig. S2A-D).

### *Prnp* loss reduces dystrophic neurites around neuritic plaques without altering Aβ accumulation

The overarching hypothesis to be tested is that PrP^C^ is required for deleterious neuronal phenotypes as a receptor, but not for the production or accumulation of Aß itself. Therefore, we measured dense core Aß plaque load and Aß accumulation with Thioflavin S and D54D2 anti-Aß antibody staining, respectively. *Prnp* knockout had no observable effect on Aß accumulation in DKI mice (Fig. 6A-C). Thus, any PrP^C^-mediated neuronal phenotypes in DKI mice occur independently of Aß accumulation, consistent with the hypothesis of PrP^C^ as a synaptic binding site for Aß oligomers.

**Figure 6.**
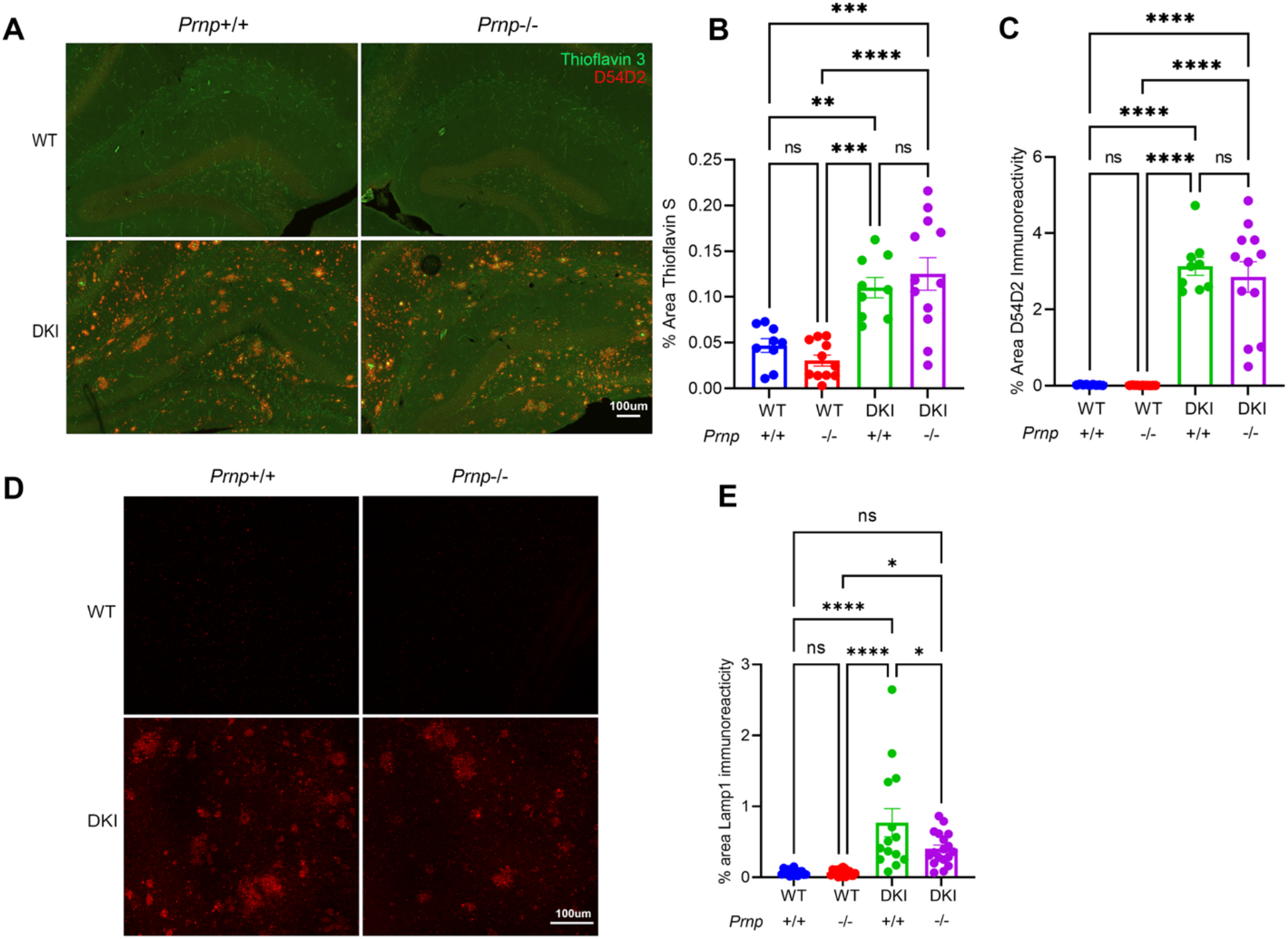
*Prnp* deletion does not alter Aß accumulation in DKI mice but reduces periplaque dystrophic neurites. (A) Representative images of Thioflavin S (green) and D54D2 anti-Aß antibody (red) staining from the hippocampus of 10-month-old WT, *Prnp^-/-^*, DKI and DKI; *Prnp^-/-^* mice. Scale bar = 100 μm. (B) Quantification of Thioflavin S stained area in hippocampus demonstrates a significant increase in dense core plaque load in DKI mice compared to WT, but no difference between DKI and DKI; *Prnp^-/-^*. Data are graphed as mean ± SEM analyzed by ordinary one-way ANOVA with Tukey’s multiple comparisons test, **P<0.01, ***P<0.001, ****P<0.0001, n= 9 for WT, n=11 for *Prnp^-/-^*, n= 9 for DKI, n= 12 for DKI; *Prnp^-/-^*. (C) Quantification of D52D2 immunoreactive area in hippocampus documents a significant increase in Aß levels in DKI mice compared to WT, but no difference between DKI and DKI; *Prnp^-/-^*. Data are graphed as mean ± SEM, analyzed by ordinary one-way ANOVA with Tukey’s multiple comparisons test, ****P<0.0001, n= 9 for WT, n= 11 for *Prnp^-/-^*, n= 9 for DKI, and n= 12 for DKI; *Prnp^-/-^*. (D) Representative images of anti-Lamp1 (red) immunostaining in medial cortex of 10-month-old WT, *Prnp^-/-^*, DKI and DKI; *Prnp^-/-^* mice. Scale bar = 100 μm. (E) Quantification of the area of Lamp1 immunoreactive accumulation in medial cortex demonstrates a significant increase of dystrophic neurites in DKI mice relative to WT. The accumulation of dystrophic neurites is less pronounced in DKI; *Prnp^-/-^* mice than in DKI mice. Data are graphed as mean ± SEM, analyzed by ordinary one-way ANOVA with Tukey’s multiple comparisons test, *P<0.05, ****P<0.0001, n= 14 for WT, n= 17 for *Prnp^-/-^*, n= 14 for DKI, and n= 18 for DKI; *Prnp^-/-^*.

The presence of Aß plaques in the AD brain is associated with local activation and increased density of microglia as well as astrocytosis. In the aged DKI brain, evidence for astrocytosis, microgliosis and microglial activation is clear (Suppl. Fig. S3). Paralleling the constant level of Aß accumulation with and without PrP^C^, the deletion of *Prnp* had no detectable effect on markers of gliosis in DKI mice.

Dystrophic neurites in the vicinity of Aß plaques are rich in lysosomal and autophagic markers and are thought to contribute to AD pathogenesis [44–46]. Therefore, we assessed whether the periplaque accumulation of neuronal organelles, which occurs in vicinity of high Aßo concentration, might require PrP^C^. We visualized dystrophic neurites by staining 10-month-old AD mouse brains with antibodies directed against lysosome-associated membrane protein-1 (LAMP-1). The DKI brain showed increased LAMP-1 immunoreactive clusters in medial cortex compared to WT mice (*p*<0.0001) consistent with a periplaque pattern, and *Prnp* knockout significantly decreased LAMP-1 accumulation on the DKI background (*p*<0.05) (Fig. 6D, E). Thus, while *Prnp* loss does not impact plaque burden or Aß staining, PrP^C^ contributes to the formation of dystrophic neurites and presumably neuronal lysosomal dysfunction.

### PrP^C^ is required for the localization of C1q to PSD-95 puncta

Microglia-mediated phagocytosis of synapses is mediated by C1q and is implicated in AD synapse loss with complement tagging synapses for removal [19, 47, 48]. We previously showed that DKI mice have increased localization of complement C1q protein to synapses compared with WT [19]. Here, we assessed the necessity of PrP^C^ for this synaptic tagging phenomenon in 10-month-old and 20-month-old mice. The DKI brain contains greater immunoblot signal for C1q than WT, but *Prnp* deletion has no impact on overall C1q levels in DKI or WT mice (Fig. 7A, B). This is consistent with the unchanged levels of gliosis and Aß accumulation. Next, we considered whether PrP^C^ alters the efficiency of C1q tagging of synapses by colocalization with PSD-95. The fraction of PSD-95 immunostaining overlapping C1q immunostaining was significantly increased in DKI mice compared to WT, while *Prnp* knockout resulted in significantly reduced PSD-95/C1q overlap on the DKI background (*p*<0.05) (Fig. 7C, D).

**Figure 7.**
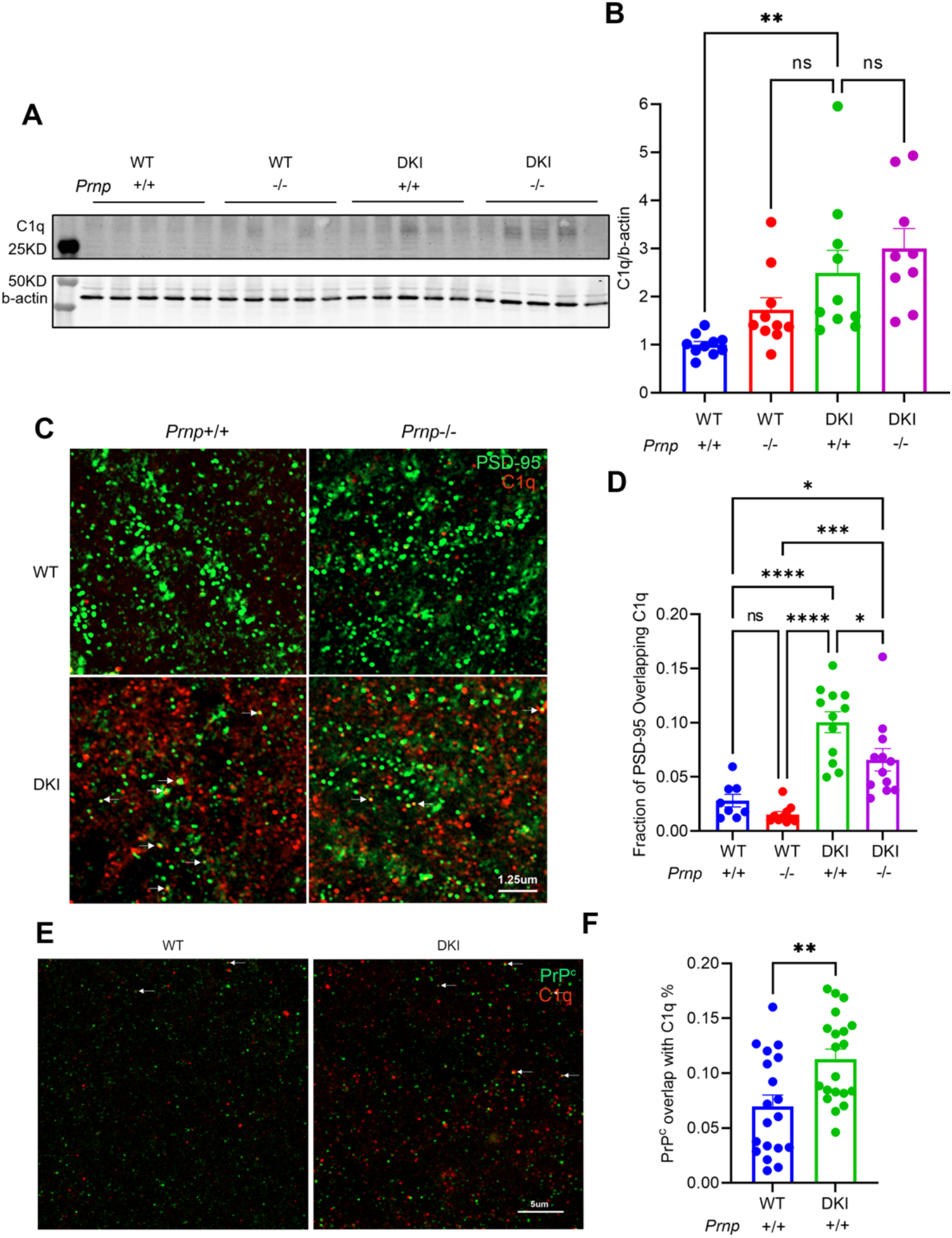
PrP^C^ mediates the localization of C1q to PSD-95 puncta without altering C1q levels. (A) Anti-C1q immunoblot image of cortical brain lysates from 20-month-old WT, *Prnp^-/-^*, DKI and DKI; *Prnp^-/-^* mice. (B) Quantification of C1q protein levels from densitometric analysis of immunoblots as in A. DKI animals exhibit significantly increased C1q expression compared to WT. The C1q level in DKI; *Prnp^-/-^* mice does not differ from DKI group values. Data are graphed as mean ± SEM, analyzed by ordinary one-way ANOVA with Dunnett’s multiple comparisons test, P>.05, **P<.01, n= 10 for WT, n= 10 for *Prnp^-/-^*, n= 10 for DKI, and n= 9 for DKI; *Prnp^-/-^*. (C) Representative images of PSD-95 (green) and C1q (red) immunoreactivity in sections of CA1 hippocampus of 10-month-old old WT, *Prnp^-/-^*, DKI and DKI; *Prnp^-/-^* mice. Scale bar = 1.25 μm. (D) Quantification of the fraction of PSD-95 immunoreactivity overlapping C1q demonstrates a significant increase in the marking of synapses by C1q in DKI animals compared to WT and WT prnp^-/-^ mice that are significantly decreased by Prnp gene knockout. Data are graphed as mean ± SEM, analyzed by ordinary one-way ANOVA with Tukey’s multiple comparisons test, *P<0.05, ***P<0.001, ****P<0.0001, n= 8 for WT, n= 10 for *Prnp^-/-^*, n= 12 for DKI, and n= 12 for DKI; *Prnp^-/-^*. (E) Representative images of PrP^C^ (green) and C1q (red) immunoreactivity in CA1 of 10-month-old WT and DKI mice. Scale bar = 5 μm. (F) Quantification of the fraction of PrP^C^ immunoreactivity overlapping C1q demonstrates a significant increase in the colocalization of PrP^C^ and C1q in DKI samples compared to WT brain. Data are graphed as mean ± SEM, analyzed by unpaired t test, **P<0.01, n= 19 for WT, n= 20 for DKI.

The reduction in synaptic tagging may reflect C1q interaction with synaptic complexes containing PrP^C^. C1q has been shown to complex with cytotoxic prion protein oligomers and to mediate the development of prion disease [49, 50]. We assessed the colocalization of PrP^C^ with C1q in 10-month-old mouse brain tissue. Immunostaining demonstrated an increase in the fraction of PrP^C^ overlapping C1q in DKI mice compared to WT (Fig. 7E, F). This data supports a potential role for PrP^C^ in the tagging of synapses by C1q, and is likely to contribute to the mechanism by which *Prnp* deletion rescues synapse loss in the DKI model.

### Transcriptomic analysis of cellular changes in DKI mice with and without PrP^C^

To provide greater insight into the cellular and molecular mechanism whereby *Prnp* deletion rescues multiple aspects of neuronal function in the DKI model, we conducted deep single-nuclei RNA sequencing (snRNA-seq) and transcriptomic analysis of WT and DKI mice with and without *Prnp* deletion at 10 and 20 months of age (Fig. 8). Cortical plus hippocampal tissue from male and female mouse brain was processed to yield 10,000 nuclei per sample, and the mean sequencing depth was 30,000 reads per nuclei (Fig 8A, B). The integrated and batch-corrected sequences were clustered in UMAP space (Fig. 8C). Cell clusters were identified by the expression of top cell-type specific classification markers (Fig. 8D). In addition to several excitatory (ExNeurons, ExN) and inhibitory neurons (InNeurons, InN) clusters, three distinct clusters of astrocyte (Astro) cell types were also identified (Fig. 8C, D).

**Figure 8.**
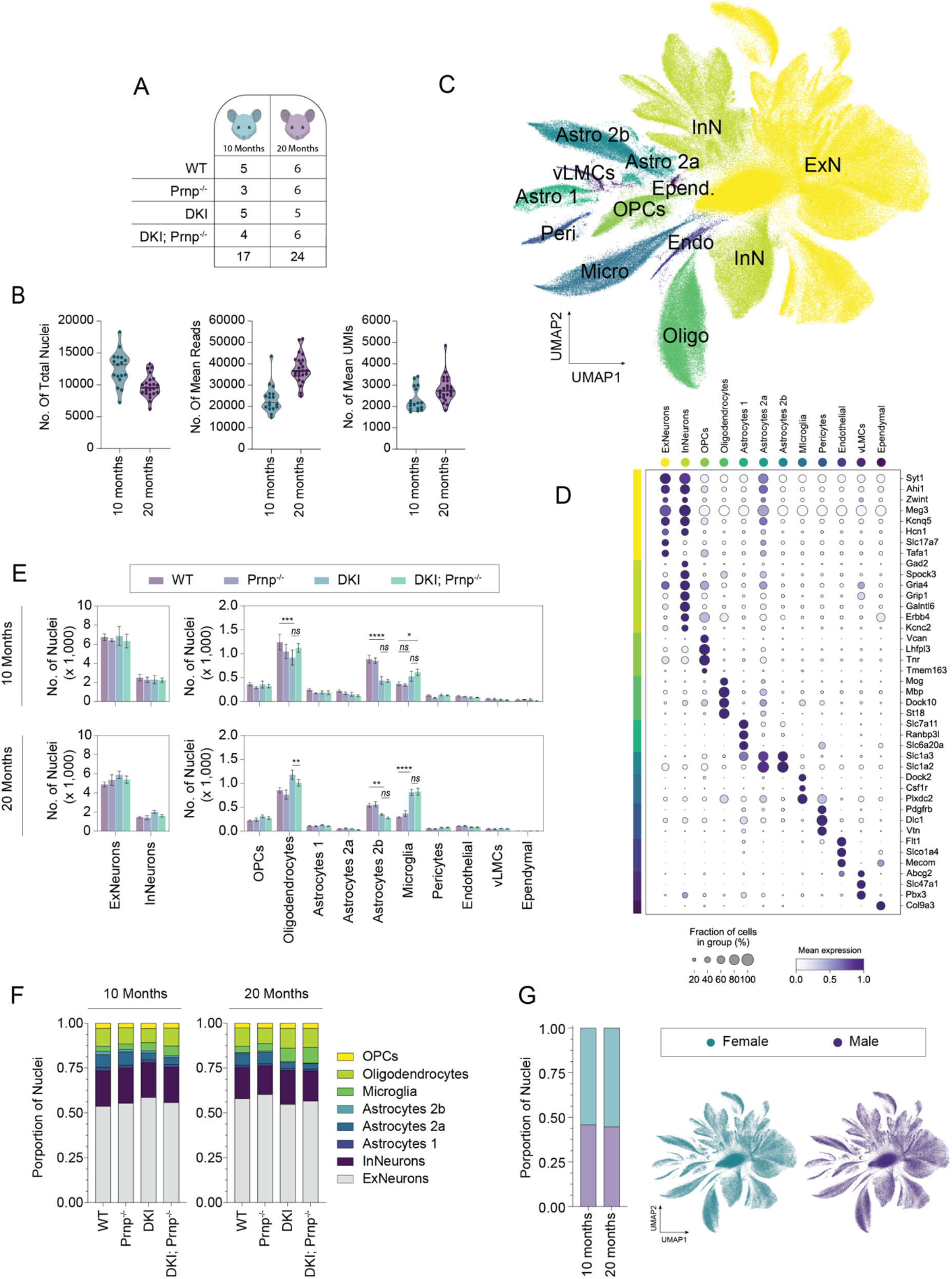
Integration and cell-type classification of age-grouped snRNA-seq data sets. (A) Schematic of treatment group and cohort sample sizes by age group. (B) Number of total nuclei per sample (left), mean reads per nuclei per sample (middle), and mean unique genes detected per nuclei per sample (right) from single-nucleus RNA-seq (snRNA-seq) of 10-month and 20-month cohorts. (D) Dot plot showing the percentage of nuclei and the scaled mean expression of cell-type classification markers used to identify the UMAP clusters in C. (E) Comparative mean nuclei frequency of cluster cell types representing individual samples of WT, *Prnp^-/-^*, DKI, or DKI; *Prnp^-/-^*genotypes. Data presented as mean+SEM from individual samples by their respective genotype and compared by mixed model test with Tukey correction. ns=non-significant, *P<0.0332, **P<0.0021, ***< 0.0002,****P<0.0001. (F) Nuclei proportions for neuronal and glial cell types across groups for 10-month and 20-month cohorts. (G) Nuclei proportions and UMAP projection of maleand female representation within the age-group integrated snRNA-seq-data.

Nuclei of all cell types were present in both age groups and sexes (Fig 8E) with male and female nuclei proportions equally represented across all cluster cell types (Fig 8G). Neuronal nuclei proportions did not vary across groups (Fig 8E-F), reflecting an absence of neuronal loss in this model. In contrast, genotype differences in glial cell proportions were observed (Fig 8E-F). At both 10 and 20 months, there was a ~50% decrease in the sub-group proportion of astrocyte 2b in DKI groups as compared with WT, with and without *Prnp*. The overall proportion of astrocyte cell types was also decreased for all genotypes at 20 months as compared to 10 months. For microglia, cell portions were higher in DKI groups as compared to WT at 10 months, with a 50% increase in microglia cell counts at 20 months. Oligodendrocyte nuclei counts were also higher in DKI groups at 20 months as compared to WT. The glial cell proportion differences occurred in DKI groups with and without *Prnp* deletion, indicating that these AD and age-dependent changes in glial number are independent of PrP^C^. This is consistent with the histological analyses described above.

Our analysis focused on transcriptomic changes within cell types, focusing first on the role of *Prnp* under non-pathological conditions (Fig. 9). *Prnp* deletion generated several differentially expressed genes (DEGs) in ExNeurons and InNeurons as compared to WT at both 10 and 20 months (Fig. 9A, B). Volcano plots revealed genes similarly up or downregulated in both ExNeurons and InNeurons (Fig. 9D), indicating a common functional role of *Prnp* in all neuronal types. While the number of neuronal DEGs remained largely unchanged between age groups, there was an age-dependent increase in *Prnp*-associated DEGs in glial (Fig 9A). Interestingly, the genetic dysregulation induced by *Prnp* deletion was greatest in oligodendrocytes, increasing 10-fold from 10 and 20 months, as compared to microglia and astrocytes (Fig. 9A,C). Of the *Prnp*-associated DEGs in WT oligodendrocytes, an adhesion molecule for neuronal cell contact, Cdh2, was most significantly downregulated (Fig 9E, see Suppl. Table S1 for full list).

**Figure 9.**
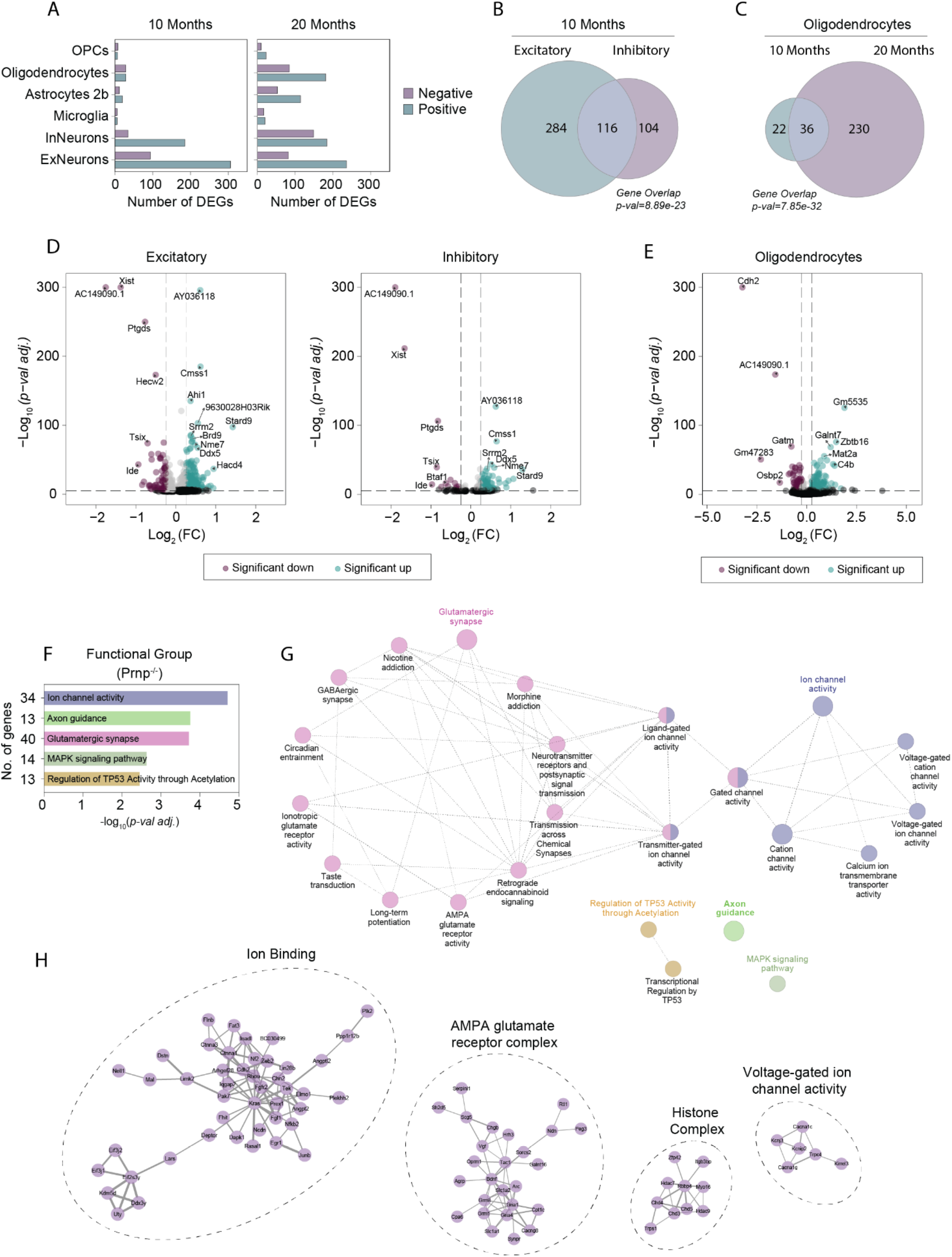
PrP^C^-dependent gene expression profile in neuronal and glial cell populations. (A) The number of positively and negatively differently expressed genes (DEGs) in neuronal and glial cell sub-populations of 10-months (left) and 20-months (right) in *Prnp^-/-^* mice as compared to WT. (B) Venn diagram demonstrating the number of shared DEGs between excitatory and inhibitory neuronal cell populations of 10-month-old *Prnp^-/-^* mice. (C) Venn diagram of significant DEGs in oligodendrocytes shared between 10- and 20-months old *Prnp^-/-^* mice. The statistical significance of gene overlap shown in B,C was assessed by Fisher’s exact test relative to the number of DEGs. (D) Volcano plots representing the gene expression changes in excitatory (left) and inhibitory (right) cells population showing the statistical significance (Log_10_(*p-val adj*.), y-axis) vs the Log2 Fold Change (Log2(FC), x-axis) of the *Prnp^-/-^* group as compared to WT; vertical dashed lines indicate ±0.25 Log2(FC). A selection of significant PrP^C^-associated genes is labeled. DEGs are deemed significant if they exhibit an absolute Log2(FC) > 0.25, with *p < 0.005* (Wilcoxon rank sum test) and are colored purple (downregulated) or teal (upregulated). For the full gene list, refer to Suppl. Table S1. (E) Volcano plot of gene expression changes identified in 20-month oligodendrocyte cell populations. (F-G) Pathway enrichment analysis of pooled significant excitatory and inhibitory DEGs between *Prnp^-/-^* versus WT mice at 10 months. (G) Enrichment terms (nodes), resulting from GO Molecular Functions, KEGG, and REAC Pathway analysis, organized into functional network groups which are linked by shared gene associations (edges). The intra-group significant terms (leading terms) are color highlighted with corresponding p-value and total functional group gene association count plotted in F. (H) Protein-Protein interaction (PPI) evidence of gene set enrichment analysis shown in F-G. Genes with stronger associations are connected by thicker lines with gene groups encircled based on shared functional pathways.

The *Prnp* deletion DEG list of 10-Month ExNeurons and InNeurons was analyzed for cellular and functional enrichment with ClueGo [34]. Gene Ontology (GO) cellular compartment analysis detected highly enriched terms related to neuronal synapses, specifically post-synaptic membrane structures (Suppl. Fig. S5, Suppl. Table S2). Gene set enrichment of functional network associations identified pathways related to glutamatergic synapses and ion channel activity (Fig. 9F-G, Suppl. Table S2), both contained strongly connected protein-protein interaction (PPI) networks involved in synaptic signal transduction (Fig. 9E). Thus, in the absence of AD-related changes, certain aspects of synaptic function are modulated by constitutive PrP^C^ loss in adult mice, and at an advanced age, oligodendrocyte transcriptomic changes are prominent in *Prnp* null mice.

### Neuronal AD-related transcriptome changes are PrP^C^-dependent while microglia are not altered

Having assessed the baseline effect of PrP^C^ loss, we turned to the DKI model and searched for AD-related transcriptomic changes dependent on *Prnp* expression (Fig. 10). Our previous work had identified numerous neuron-specific DEGs rescued to a normal expression pattern by treatment with BMS-984923, a silent allosteric modulator of mGluR5 [19]. Based on PrP^C^ interaction with mGluR5 [21, 51], we hypothesized a related pattern here. Consistent with our previous findings and other studies, we observed numerous AD-associated DEGs in DKI compared to WT. At 10 months during the early phenotypic stage, the number of DEGs are considerably higher in neurons as compared to glial cell types (Fig. 10A, Suppl Table S1). Conversely, AD-associated gene expression changes in glial cells were more numerous at 20 months during disease progression as compared to 10 months (Suppl. Fig. S4A, B). In neurons, the number of DEGs was greater in excitatory than inhibitory neurons, though many DEGs were shared between excitatory and inhibitory neurons (Fig. 10B, C). The list of neuronal DEGs included upregulated *Camk2n1, Grin2b*, and *Chchd3*, genes essential for proper synaptic maintenance and stability, and for which dysregulation has been associated with AD pathology. Of note, neuronal DEGS are more numerous at the 10-month phenotypic initiation timepoint than at 20 months (Fig 10C). Thus, this DKI model is characterized by early changes across neuronal cell types and delayed changes in multiple glial cell types.

**Figure 10.**
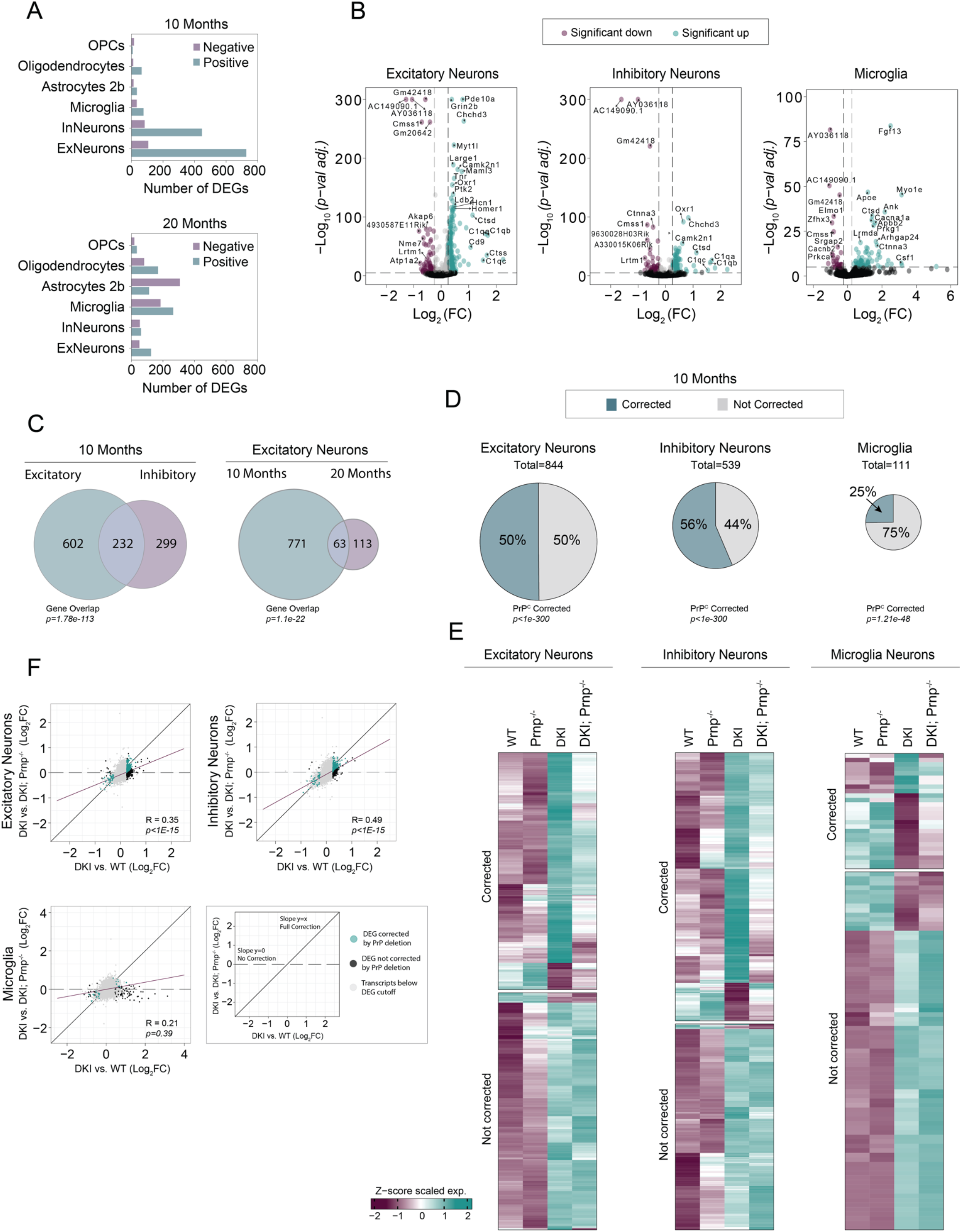
*Prnp*-dependence of cell type specific gene expression in DKI mouse model. (A) The number of AD-associated DEGS in neuronal and glial cell population at 10 (top) and 20-months (bottom) old DKI mice. (B) Volcano plots of excitatory and inhibitory neurons, and microglia at 10 months. Colored points indicated significant up or downregulated genes in DKI samples as compared to WT. (C) Venn diagrams showing the number of shared DEGs between excitatory and inhibitory neurons at 10 months, excitatory neurons populations at 10 and 20 months. The statistical significance of gene overlap was assessed by Fisher’s Exact test. (D) Pie charts illustrating the percentage of DEGs that are fully corrected by PrP^C^ gene deletion (Fisher’s exact test); the size of the chart is relative to total number of DEGs. (E) Heatmaps showing single DEG expression within each cell type, separated by corrected and non-corrected genes as represented in D,F. DEG lists showing the remaining cell types are included in Supp. Table S2. (F) Transcriptome-wide correction by PrP^C^ gene deletion in neuronal and glial cell types. Cell type-specific comparison of AD-associated DEGs and PrP^C^-corrected DEGs in DKI samples. Log2FC between DKI and WT samples (AD effect) is plotted along the *x-*axis. Log2FC between DKI and DKI; *Prnp^-/-^* samples (PrP^C^ effect) is plotted along the *y-axis*. Black points represent genes with Log2FC > 0.25 and *p < 0.005*. Colored points represent DEGs that were corrected by PrP^C^ deletion. Points along the identity line (*x* = *y*) represent genes with equivalent differential expression between DKI; *Prnp^-/-^* and WT, relative to DKI, indicating complete rescue by PrP^C^ deletion. Points along the line *“y* = 0” reflect genes unaffected by PrP^C^. *P*-values represents the significance of a non-zero linear regression relationship. The regression line (Pearson’s correlations, purple) represents transcriptome-wide effects of PrP^C^.

Having established a temporal profile for transcriptomic response to the AD-related gene knock-ins, we examined whether *Prnp* deletion would correct the dysregulation in DKI; *Prnp^-/-^* neurons, normalizing expression levels to WT levels. Indeed, for excitatory and inhibitory neurons there was a 50% and 56% correction of DEGs, respectively, in response to *Prnp*-deletion (Fig. 10D-E). In contrast, only 25% of genes were corrected by *Prnp* deletion in microglia at 10 months. To understand the transcriptomic-wide effect of *Prnp* deletion in our DKI model of AD, we plotted the gene Log2 fold-change (Log2FC) of DKI vs WT (AD-effect, *x-axis*) against that of DKI vs DKi; *Prnp^-/-^* (deletion effect, *y-axis*). Using linear regression analysis on all plotted genes, a linear regression with slope=1 represented full correction, and with slope=0 corresponded to no correction of the overall AD-associated transcriptomic profile (Fig. 10F). Both excitatory and inhibitory neurons had a higher slope of regression (purple line) with corrected genes (teal) falling on the regression line of full correction (black). The linear regression was significantly different from a slope = 0 (*p<1E-15*) in both neuron types, again indicating a similar functional effect of *Prnp* across neuronal cell populations. In contrast, for microglia, the regression line slope was shallow and not significantly different from 0 (p=0.39). Thus, *Prnp* has little transcriptomic effect on microglia cells at 10 months (Fig. 10F). To characterize the role of *Prnp* in DKI neurons, we performed functional and pathway analyses using the pooled gene set list of *Prnp*-corrected neuronal DEGs. Top functionally enriched networks include the cGMP-PKG, WNT, and RAS/MAPK pathways (Fig. 11A-C, Suppl Table S2). These are all second-messenger signaling pathways involved in the modulation of synaptic transmission and long-term potentiation. Thus, *Prnp* is required for early AD-related disruption of these neuronal pathways.

**Figure 11.**
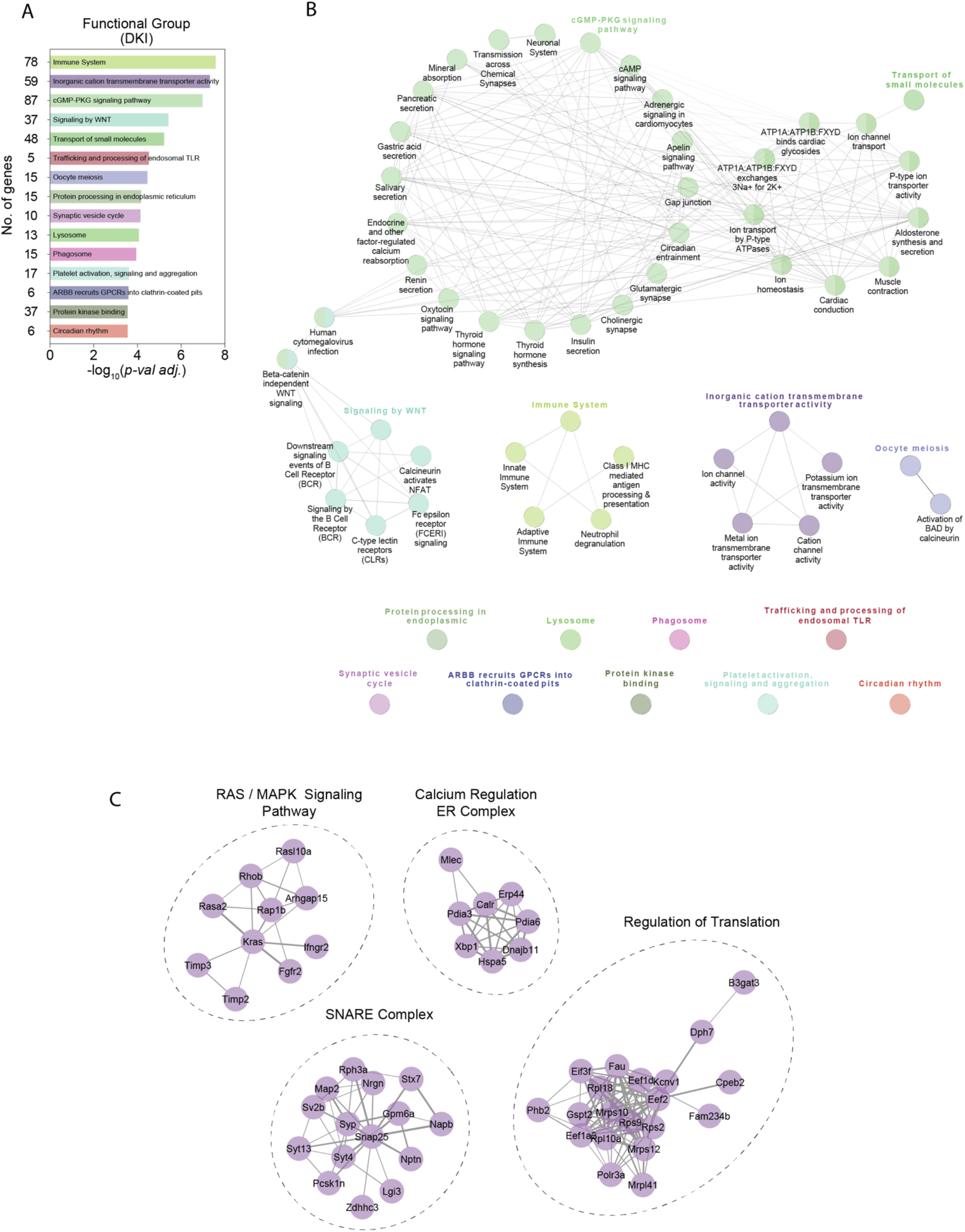
Pathways expression-normalized in DKI neurons by loss of PrP^C^ function. Functional enrichment analysis of AD-associated DEGs (DKI versus WT) corrected by *Prnp* deletion in pooled excitatory and inhibitory neuron samples. (A) Leading terms are color highlighted with corresponding p-values and total gene association count of each function group is graphed. (B) Gene set enrichment analysis against GO Molecular Function terms, KEGG and REAC Pathways, organized into functional network groups (nodes) based on shared gene associations (edges). (C) Functionally enriched PPI networks (encircled) with genes having stronger associations connected by thicker lines and genes with lesser or unknown associations not shown.

With regard to glial cells, the number of DEGs in DKI as compared to WT is higher at 20 months (Fig. 10A). We considered whether the temporal differences were related to a simple delayed onset or a shift in the expression pattern of glial reaction. We assessed whether gene expression changes at 20 months were also present at 10 months (Suppl. Fig. S4). In general, top DKI glial cell DEGs were uniquely present at 20 months, with little to no altered expression at 10 months (Fig. S4A). The notable exception was in microglia, for which most AD-associated DEGs had similar expression patterns at both 10 and 20 months (Suppl. Fig. S4B). In this regard, microglia are part of the disease initiation phenotype, while astrocytic and oligodendrocyte populations participate more specifically in later disease progression. Interestingly, lipoprotein genes *Apoe* and *Apod*, proteolytic protein genes *Ctsb, Ctsd*, and *Ctsl*, as well as complement gene *C1q* are also differentially expressed in both neurons and oligodendrocytes at 10 months.

Transcriptomic changes occur earlier and are stronger in microglia than in oligodendrocytes and astrocytes (Suppl. Fig. S4B, D). However, the functional effect of *Prnp* differed between glial cell types. At 20 months, we found that 26% of oligodendrocyte and 32% of astrocyte DEGs were corrected by *Prnp* deletion (Suppl. Fig. S4C). However, *Prnp* deletion only corrected 10% of AD-associated microglia DEGs. Transcriptomic-wide profile analysis of microglia had a linear regression slope of 0 (p=0.39), indicating that *Prnp* has no direct role functional role in microglia (Suppl. Fig. S4D). Conversely, OPCs, oligodendrocytes, and astrocytes had transcriptomic profiles with linear regressions significantly different from a slope = 0. Although, there was the normalization of some AD-associated DEGs in oligodendrocytes and astrocytes cell populations of DKI; *Prnp^-/-^* mice at 20 months, there was also genetic dysregulation with *Prnp* deletion alone when compared to WT (Fig 8A). This is also clear in Venn diagrams of shared and unique DEGs between *Prnp^-/-^*, DKI, and DKI; *Prnp^-/-^* as compared to WT (Suppl. Fig. S4E). Therefore, PrP^C^ appears to play a maintenance role in oligodendrocyte and astrocyte cell populations, as well as contributing to delayed and potentially indirect AD-related PrP^C^ signaling.

While investigating to what extent *Prnp* deletion could normalize AD-associated gene expression changes, we also found several dysregulated genes unique to the DKI; *Prnp^-/-^* genotype (Fig. 12A). This DEG set is dependent on the synthetic interaction of *Prnp* loss with the AD model and is not detected with either single condition. The observed synthetic interaction was specific to neurons in the 10-month DKI; *Prnp^-/-^* brain (Fig. 12B-F). The phenomenon was not observed in glial cells (Fig. 12B, Suppl. S4E). Specifically, the number of DEGs for each of the DKI; *Prnp^-/-^* glial cell types were at levels similar to *Prnp^-/-^* with many DEGs shared between the two genotypes (Suppl. Fig S4E). The 10-month neuronal synthetic interaction DEG set may reflect compensatory signaling mechanisms in DKI; *Prnp^-/-^* neurons, so we performed gene ontology pathway enrichment analysis (Suppl. Fig S6, Suppl Table S2). Within the synthetic interaction set, nearly all enriched cellular compartment GO terms were associated with post-synaptic localization (Suppl. Fig. S5). With regard to functional pathway enrichment, the same glutamatergic and ionic conductance pathways associated with *Prnp^-/-^* neurons were observed, but the top significant enrichment terms unique to the synthetic interaction set included Rho GTPase and cytoskeleton activity (Suppl. Fig. S6A-B). These Rho and cytoskeletal enrichment terms also had the most highly connected gene-network associations (Suppl. Fig. S6C). Collectively, our transcriptomic profiling results show that *Prnp* deletion normalizes DKI-dependent neuronal expression patterns specifically related to synaptic function and stability both by simple corrective rescue and by compensatory mechanisms.

**Figure 12.**
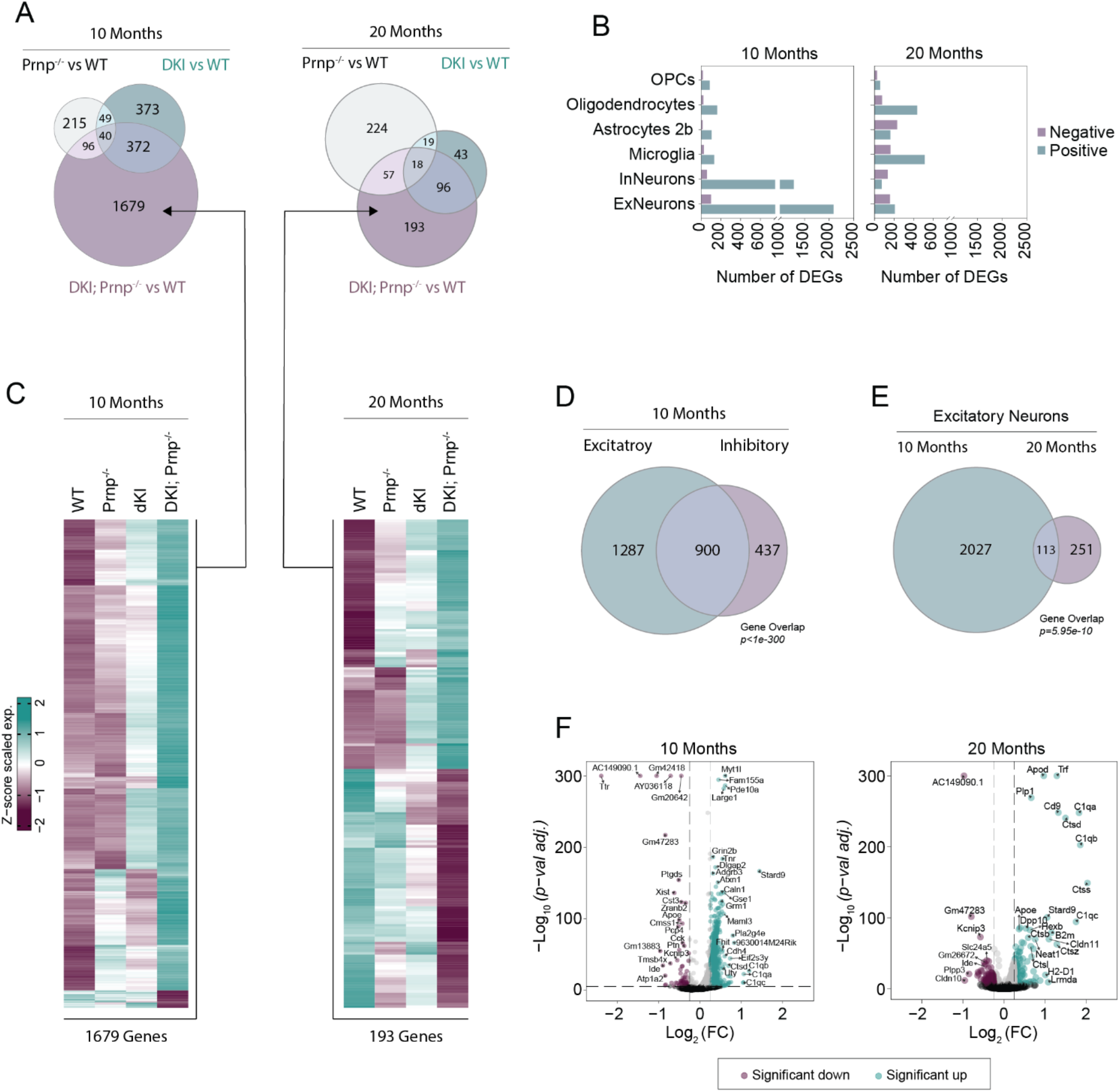
Synthetic gene expression profiles dependent on the interaction of DKI model with *Prnp* deletion. (A) Venn diagrams depicting the number of shared and unique DEGs between *Prnp^-/-^*, DKI, and DKI; *Prnp^-/-^* samples in 10 (left) and 20 month (right) excitatory neuronal cell populations. (B) The number of positive and negative DEGs detected within neuronal and glial cell populations DKI; *Prnp^-/-^* mice at 10 (left) and 20 months (right). (C) Heatmaps showing single DEG expressions of unique to DKI; *Prnp^-/-^* samples (arrows) as represented in A. (D,E) Venn diagrams demonstrating the shared DEG significance (Fisher’s Exact Test) between excitatory and inhibitory neuronal cell population at 10 months (D) and between cell populations of excitatory neurons at 10 and 20 months. (F) Volcano plots showing DEGs of 10 (left) and 20 month (right) excitatory neuronal cell populations as measured by snRNA-seq comparing DKI; *Prnp^-/-^* to WT samples.

## DISCUSSION

The primary finding of the current study is that PrP^C^ mediates multiple neuronal AD-related phenotypes in mice with endogenous expression at the *Mapt* and *App* loci (*App^NL-G-F^/hMapt* double knock-in, or DKI). DKI mice demonstrate spatial memory deficits at 9 months that require a functional *Prnp* gene. Synapse loss, as measured by immunohistochemistry and SV2A PET, is also fully rescued by *Prnp* gene deletion and coincides with a significant decrease in synaptic tagging by C1q in DKI; *Prnp^-/-^* mice. DKI mice demonstrate appreciable phospho-tau accumulation that is significantly reduced in the absence of PrP^C^. The effects of *Prnp* gene deletion appear to be primarily neuronal, as evidenced by significant correction in DKI-dependent neuronal gene expression changes with *Prnp* gene deletion. In stark contrast, glial gene expression changes in DKI mice are only minimally altered by *Prnp* gene deletion, and microgliosis, astrocytosis, and Aß levels are unchanged between DKI and DKI; *Prnp^-/-^* mice. Several biomarkers of the essential role of PrP^C^ in neuronal AD phenotypes are translatable to clinical scenarios, including SV2A PET [27] and pThr217 levels [52], and these may be used to assess the therapeutic efficacy of targeting Aßo-PrP^C^ interaction in AD.

Neuronal metabotropic glutamate receptor 5 (mGluR5) is a transmembrane protein whose aberrant activation links the extracellular Aßo-PrP^C^ interaction to intraneuronal signaling in mouse models of AD [10, 18, 20, 21]. We recently demonstrated the efficacy of a silent allosteric modulator (SAM) of mGluR5 signaling to rescue AD phenotypes in the DKI model [19]. Similar to *Prnp* gene deletion, SAM administration rescued synapse loss, reduced phospho-tau accumulation, and reduced synaptic tagging with C1q in DKI mice. Additionally, SAM corrected DKI-dependent genes primarily in neurons with little change observed in glial cells through transcriptomics and immunohistochemistry. The mirroring of SAM and *Prnp* deletion in rescuing DKI mice is consistent with their co-joined function as a molecular complex [21, 51, 53]. These data demonstrate that disrupting the Aßo-PrP^C^-mGluR5 pathway at two different steps rescues AD phenotypes, primarily through neuronal expression changes and with modulation of neuronal-glial interactions via C1q. Despite the convincing similarity in mechanistic rescue of AD phenotypes with SAM administration and *Prnp* deletion, there are important differences in these two studies. The *Prnp* deletion study utilizes a constitutive knockout in a prophylactic mode prior to any AD related pathology, while the SAM treatment was effective in a therapeutic mode after the DKI mice were aged to the point of synapse loss (~12 month development here). Additionally, *Prnp*-null mice show transcriptomic changes separate from DKI effects, consistent with PrP^C^ having many purported physiological functions, including regulation of myelin maintenance, cell differentiation, and neuronal excitability [54]. In contrast, pharmacological targeting of mGluR5 with a SAM has minimal if any effect in WT mice [19, 53]. It remains possible that the perturbation of one of the many roles of PrP^C^ other than Aßo binding might contribute to the rescue of AD phenotypes in the DKI model. However, the well-described interaction of PrP^C^ with mGluR5 and mechanistic congruency with pharmacologically targeting mGluR5 suggest that the therapeutic effects of *Prnp* deletion on AD pathology are largely mediated through the Aßo-PrP^C^-mGluR5 pathway.

We observed that deletion of *Prnp* normalizes DKI-dependent neuronal expression patterns, especially in synaptic networks, but with little change in glial populations. Similar to SAM administration, in which only 7% of microglia DEGs were corrected with drug administration [19], *Prnp* deletion corrected only 25% of microglia DEGs in DKI mice. Meanwhile, *Prnp* deletion corrected greater than half of DEGs in excitatory and inhibitory neurons. As is consistent with previous studies investigating disruption of the Aßo-PrP^C^-mGluR5 pathway [14, 15, 19, 20], overall disease-associated changes in microgliosis, astrocytosis, microglial activation, total C1q levels, plaque load, and overall Aß levels are unchanged. Despite glial gene expression and overall levels being largely unchanged, their interaction with neurons appears to be modulated through PrP^C^-dependent transcriptomic changes and tagging of synapses by C1q. Thus, targeting PrP^C^ or mGluR5 pharmacologically rescues AD phenotypes through neuronal gene expression changes that abrogate the aberrant deleterious interactions between glia and neurons. Further investigations are necessary to understand these possible mechanisms, including specific C1q receptors and the molecular role of PrP^C^ in the localization of C1q to synapses and subsequent synaptic phagocytosis by microglia and/or astrocytes.

*Prnp*-null mice exhibit phenotypes that underscore the broad and still partially understood physiological functions of PrP^C^. At 10 months age, mice with a targeted *Prnp* deletion demonstrated significantly reduced [^18^F]SynVesT-1 uptake in the olfactory bulb and increased uptake in the caudate-putamen. The importance of PrP^C^ in maintaining mature olfactory sensory neurons has been reported [38–40], and the loss of synapses as measured by SV2A PET further supports a function of PrP^C^ in proper olfactory tract development. The physiological role of *Prnp*-null specific synaptic density increases observed in caudate-putamen is not clear and merits future study. We did not conduct detailed assessment of motor function here. However, we observed no difference in swim speed or learning deficits between WT and *Prnp^-/-^* mice. The rescue of DKI phenotypes by *Prnp* deletion appears specific to the context of pathologic Aß and Tau.

Transcriptomic changes in *Prnp*-null mice were found primarily in excitatory and inhibitory neurons at 10 months age, with differentially expressed genes clustering in synaptic glutamate and ionic conductance pathways. PrP^C^ has been shown previously to have neuronal excitability-modifying properties [55], and Aß has been shown to require PrP^C^ to inhibit ionic conductance [56]. PrP^C^ has also been shown to play a role in Ca^2+^ signaling through interactions with glutamate receptors [21, 57]. These studies are consistent with our observation of PrP^C^-dependent transcriptomic changes. However, distinct neuronal expression occurred in the DKI mice and their rescue by *Prnp* deletion was largely non-overlapping with PrP^C^-dependent changes on the WT background.

By 20 months age, expression of PrP^C^ altered oligodendrocyte transcriptomic patterns. PrP^C^ has been shown to play a role in oligodendrocyte differentiation and development, with oligodendrocyte precursor cells lacking PrP^C^ proliferating more vigorously and at the expense of differentiation [58]. Our results support a role for PrP^C^ in maintaining proper oligodendrocyte function, though this was not detectable until middle age. The difference between PrP^C^-dependent oligodendrocyte changes in 10-month-old mice compared to 20-month-old mice is dramatic. While we also observed oligodendrocyte DEGs in DKI mice at 10 months and 20 months, the extent was far less than PrP^C^-dependent oligodendrocyte DEGs. Additionally, DKI-dependent DEGs occur more widely in cells like astrocytes and microglia at 20 months, while PrP^C^-dependent DEGs predominate in oligodendrocytes at 20 months.

Interestingly, a large group of neuronal DEGs were detected only as a synthetic phenotype of the knock-in/knock-out interaction of DKI; *Prnp^-/-^* mice. Since behavioral and histological phenotypes are reduced in this condition, the synthetic interaction DEG subset may have a compensatory and beneficial effect to mitigate DKI-dependent changes and restore normal physiological function. At 10 months age, these genes were primarily observed in excitatory and inhibitory neurons, while at 20 months, microglia and oligodendrocytes showed the highest presence of synthetic interaction DEGs. Network analysis reveals that they are found in a variety of pathways not typically associated with disease pathogenesis. These include ribonucleotide binding, GTPase activator activity, and signaling by Rho GTPases. Further investigation into these genes, including a comparison with DKI mice with late conditional *Prnp* deletion or pharmacological inhibition of Aßo-PrP^C^ binding, should help elucidate their function in disease pathology and treatment. Notably, such synthetic interaction DEGs were not observed with mGluR5 SAM rescue of DKI mice [19].

Taken together, these data support the hypothesis that PrP^C^ may be a safe and viable target for pharmacological intervention in AD treatment. In fact, several potential therapeutics focused on disrupting the Aßo-PrP^C^-mGluR5 pathway have been shown to be preclinically efficacious. Anti-PrP^C^ antibody that crosses the blood-brain barrier and competitively inhibits Aßo oligomer binding has been shown to rescue impaired LTP, behavioral deficits, and synapse loss [14, 59, 60]. Similarly, a polymer that binds the Aßo-binding site on PrP^C^ rescues several AD phenotypes and prevents hydrogel formation [15, 61]. mGluR5 has also been targeted through a silent allosteric modulator (SAM) which has shown strong preclinical promise and is currently in phase I clinical trials [19]. Methods described in this paper, including SV2A PET and phospho-tau profiling, can be used to evaluate the effects of these drugs in human subjects and understand any differences between human AD and mouse models.

Although endogenous expression at the *Mapt* and *App* loci potentially serves as a more accurate reflection of human AD progression than overexpressing Aß models, there are limitations to our study design. The utilization of WT mice (*App^+/+^, Mapt^+/+^, Prnp^+/+^*) as a control group does not eliminate the possibility that *hMapt* contributes to observed phospho-tau accumulation independently of Aß. It has been demonstrated that *hMapt* knock-in mice propagate pathological Tau species quicker than WT mice after injection with AD-derived tau [25]. However, in this same study, DKI mice also showed significantly increased phospho-tau accumulation compared to *hMapt* knock-in mice, suggesting at minimum an exacerbating role of Aß in the formation of tau aggregates. Importantly, humanization of the *Mapt* gene does affect memory, neuroinflammation, and Aß40 and Aß42 levels on a wild type or *App^NL-G-F^* background, supporting the hypothesis that Aß production acts upstream to initiate Tau hyperphosphorylation. Our study aimed to specifically investigate PrP^C^-dependence in this double knock-in model rather than the changes individually associated with *hMapt* and *App*^NL-G-F^ knock-in. Here, the reduction in pathological tau aggregates with *Prnp* gene deletion, a high-affinity binding partner of oligomeric Aß, supports our hypothesis that *Prnp* facilitates downstream neurotoxicity, including phospho-tau accumulation. Nonetheless, future studies comparing *App^+/+^, Mapt^hMapt/hMapt^, Prnp^+/+^* mice to *App^NL-G-F/NL-G-F^, Mapt^hMapt/hMapt^, Prnp^+/+^* mice may help to isolate the initiating effects of Aß in this model as well as understand the extent to which other mechanisms or Aß binding partners contribute to tau pathology.

## CONCLUSIONS

PrP^C^ is a high-affinity, oligomer-specific Aß binding protein required for memory loss, synapse deficits, and selective neuronal degeneration in transgenic mice overexpressing Aß. Our previous work has investigated and demonstrated the efficacy of targeting the Aßo-PrP^C^ interaction with anti-PrP^C^ antibody blockade, PrP antagonists, and gene knockout in mouse models overexpressing Aß. Here, we show that a functional *Prnp* gene is required for behavioral deficits and synapse loss in an AD mouse model with endogenous expression at the *App* and *Mapt* loci. Phospho-tau accumulation, a pathologic hallmark of AD that is limited or absent in other Aß models, is detected in the DKI model and significantly reduced with targeted *Prnp* gene deletion. While single-nuclei transcriptomics reveals differential expression in neurons and glia of DKI mice relative to WT, glial DKI-dependent gene expression changes largely persist with *Prnp* deletion, consistent with unchanged histologic levels of microgliosis, astrogliosis, and Aß levels between DKI and DKI; *Prnp^-/-^* mice. DKI-dependent neuronal gene expression changes are significantly corrected with *Prnp* deletion, and corrected genes are primarily associated with synaptic function. These data support the efficacy of targeting the Aßo-PrP^C^ interaction to prevent Aßo-neurotoxicity and pathologic tau accumulation in AD. The mechanisms by which PrP^C^ mediates AD-related phenotypes, including synaptic tagging by C1q, formation of dystrophic neurites, and phospho-tau accumulation, require further investigation.

## ABBREVIATIONS

Astro: Astrocytes
cGMP: cyclic guanosine monophosphate
DEGs: Differentially expressed genes
DKI: Double Knock-In, *App^NL-G-F/NL-G-F^, Mapt^hMAPT/hMAPT^, Prnp^+/+^*
DKI; *Prnp^-/-^*: *App^NL-G-F/NL-G-F^*, *Mapt^hMAPT/hMAPT^, Prnp^-/-^*
Epend: Ependymal
ExN: Excitatory neurons
FC: Fold change
GO: Gene Ontology
InN: Inhibitory neurons
MCL: Markov Cluster Algorithm
Micro: Microglia
Peri: Pericytes
PKGs: cGMP-dependent protein kinases
PPI: Protein-Protein Interactions
*Prnp^-/-^*: *App^+/+^, Mapt^+/+^, Prnp^-/-^*
Oligo: Oligodendrocytes
OPCs: Oligodendrocytes precursor cells
snRNA-seq: Single-nucleus RNA-sequencing
UMAP: Uniform manifold approximation projection
vLMCs: Vascular Leptomeningeal Cells
WT: Wild-type, *App^+/+^, Mapt^+/+^, Prnp^+/+^*

## DECLARATIONS

### Ethics approval and consent to participate

Not applicable.

### Consent for publication

All authors have approved of the consents of this manuscript and provided consent for publication.

### Availability of data and materials

Raw FASTQ sequencing files and gene count matrices have been deposited in NCBI’s Gene Expression Omnibus (GEO) under accession code GSE221712.

### Authors’ contributions

Conceptualization, A.S. and S.M.S.; Methodology, A.S., L.F., L.N., J.S., C.Z., T.T., Z.C., and S.M.S.; Investigation, A.S., L.F., L.N., J.S., T.T., and W.L.; Writing – Original Draft, A.S., L.F., L.N. and S.M.S.; Writing – Review & Editing, all; Funding Acquisition, S.M.S.; Resources, S.M.S; Supervision, S.M.S. All authors read and approved the final manuscript.

## Acknowledgements

We thank Le Zhang for helpful discussions and Kristin Deluca for expert technical assistance.

## FIGURE LEGENDS

### ADDITIONAL INFORMATION

Supplemental Tables S1-S2 Description and Supplemental Figures S1-S6.

**Supplemental Tables S1. List of Differentially Expressed Genes.**

Data from the experiment represented in main Figure 8 are presented. The data are separated by genotype, cell type and age. Genes meeting the defined threshold for differential expression relative to WT group of the same age, their respective Z-score, Log2 Fold Change relative to WT, and the adjusted p-value are shown. Provided as Excel table.

**Supplemental Tables S2. Gene Set Enrichments for Differentially Expressed Gene lists.**

The neuronal DEGs listed in Suppl Table S1 were assessed for cellular compartment and functional pathway Gene Set enrichment using ClueGo. Three separate DEG lists were analyzed. The *“Prnp* Null” model utilized the DEG from *Prnp^+/+^* versus WT. The *“Prnp* corrected” model utilized those genes present the DEG list from DKI versus WT, but not in the DKI; *Prnp^-/-^* versus WT list. The *“Prnp* interaction DKI” model utilized DEGs found only in the DKI; *Prnp^-/-^* versus WT list. Separate tabs are provided for each model by Functional Pathway analysis and one tab for the cellular compartment analysis of all three models. Provided as Excel table.

**Supplemental Figure S1.**
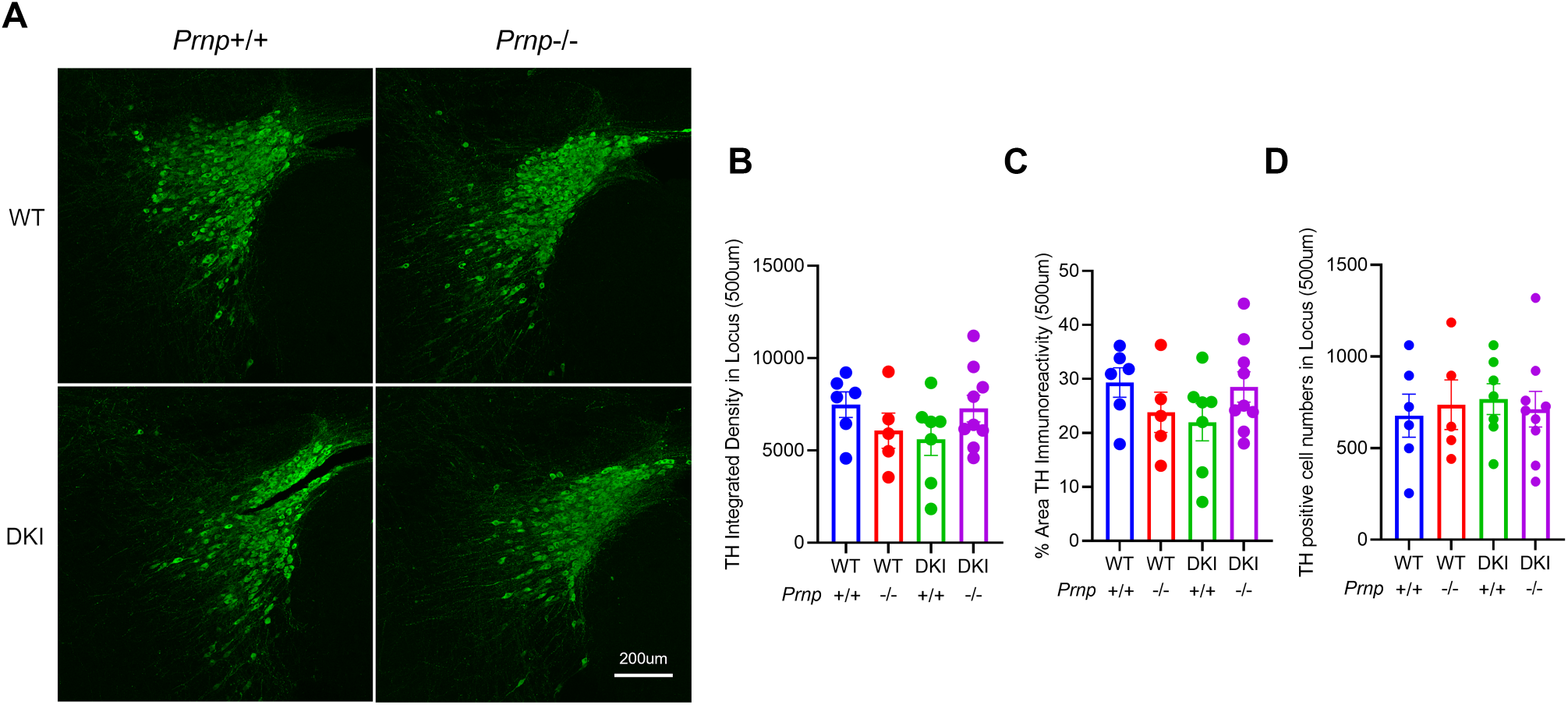
Limited locus coeruleus neuron loss in DKI mice. (A) Representative immunofluorescent images of anti-Tyrosine hydroxylase (TH) staining in locus coeruleus of 20-month-old WT, *Prnp^-/-^*, DKI and DKI; *Prnp^-/-^* animals. Scale bar = 200 μm. (B-D) Quantification of locus coeruleus TH immunoreactivity by evaluating TH integrated density (B), percentage of TH immunoreactive area (C), and TH positive cell numbers in locus coeruleus (D). There is a non-significant trend to decrease TH integrated density and TH immunoreactive area in DKI relative to WT. There is no numerical difference between WT and DKI; *Prnp^-/-^*. Data are graphed as mean ± SEM, analyzed by ordinary one-way ANOVA with Dunnett’s multiple comparisons test, P> 0.05, n=6 for WT, n=5 for *Prnp^-/-^*, n=7 for DKI, and n=9 for DKI; *Prnp^-/-^*.

**Supplemental Figure S2.**
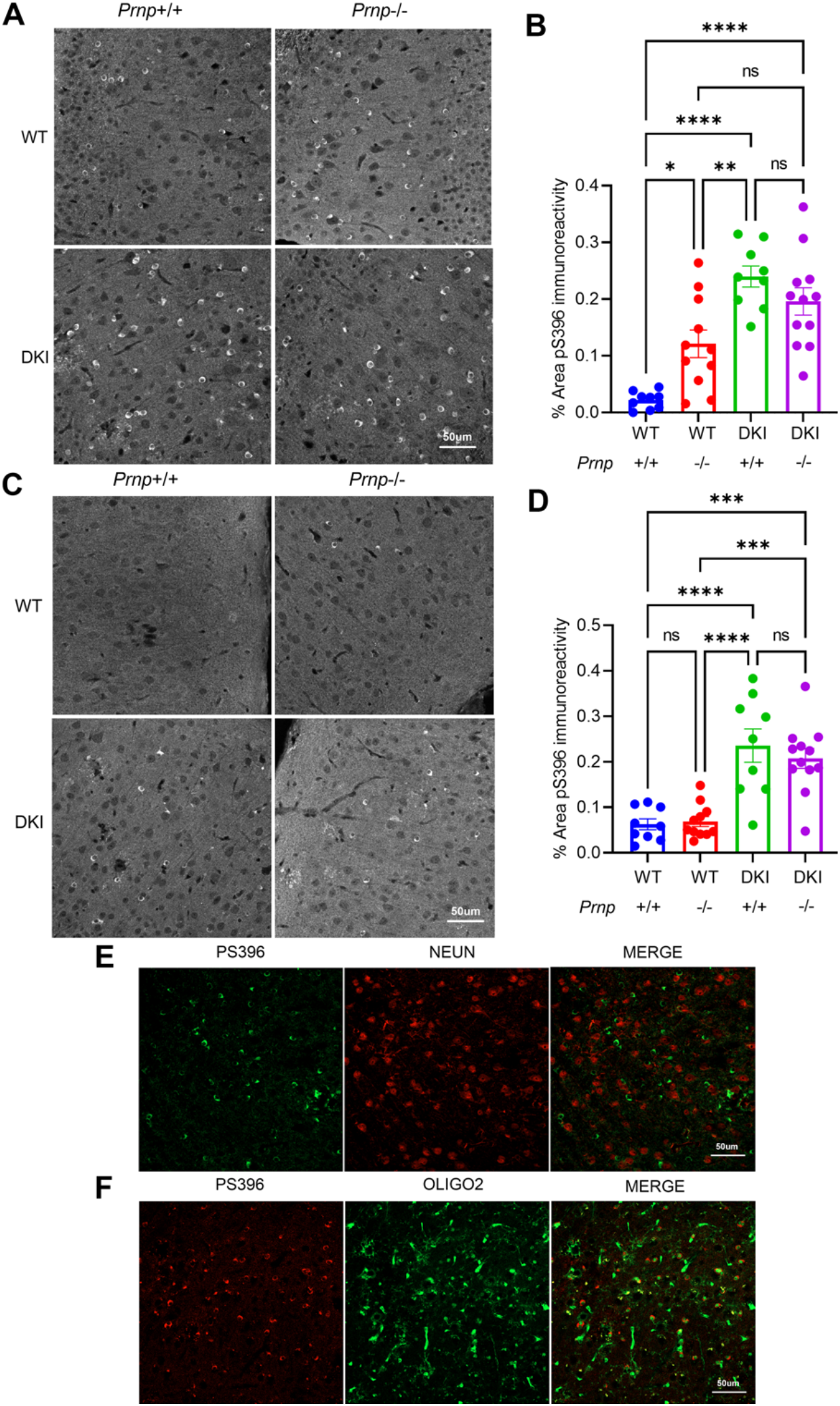
Phospho-S396 tau accumulation in DKI mice localized to oligodendrocytes. (A) Representative images of anti-phospho-S396 tau immunostaining from the medial cortex of 10-month-old WT, *Prnp^-/-^*, DKI and DKI; *Prnp^-/-^* mice. Scale bar = 50 μm. (B) Quantification of pS396 immunoreactive area from sections as in A demonstrates a significant increase for pS396-tau accumulation in *Prnp^-/-^*, DKI and DKI; *Prnp^-/-^* brain relative to WT. The DKI and DKI; *Prnp^-/-^* levels are indistinguishable. Data are graphed as mean ± SEM, analyzed by ordinary one-way ANOVA with Tukey’s multiple comparisons test, *P<0.05,**P<0.01, ****P<0.0001, n=9 for WT, n=11 for *Prnp^-/-^*, n=9 for DKI, and n=12 for DKI; *Prnp^-/-^*. (C) Representative images of anti-phospho-S396 tau immunostaining from the lateral cortex of 10-month-old WT, *Prnp^-/-^*, DKI and DKI; *Prnp^-/-^* mice. Scale bar = 50 μm. (D) Quantification of pS396 immunoreactive area from sections as in C demonstrates a significant increase for pS396-tau accumulation in *Prnp^-/-^*, DKI and DKI; *Prnp^-/-^* brain relative to WT. The DKI and DKI; *Prnp^-/-^* levels are indistinguishable. Data are graphed as mean ± SEM, analyzed by ordinary one-way ANOVA with Tukey’s multiple comparisons test, ***P<0.001, ****P<0.0001, n=9 for WT, n=11 for *Prnp^-/-^*, n=9 for DKI, and n=12 for DKI; *Prnp^-/-^*. (E) Representative immunostained images of pS396 (green) and NeuN (red) taken in the medial cortex of 10-month-old DKI mice. pS396 distribution in DKI mice is not localized to neurons. Scale bar = 50 μm. (F) Representative immunostained images of pS396 (red) and Olig2 (green) taken in the medial cortex of 10-month-old DKI mice. The pS396 signal in DKI mice is localized almost exclusively to oligodendrocytes. Scale bar = 50 μm.

**Supplemental Figure S3.**
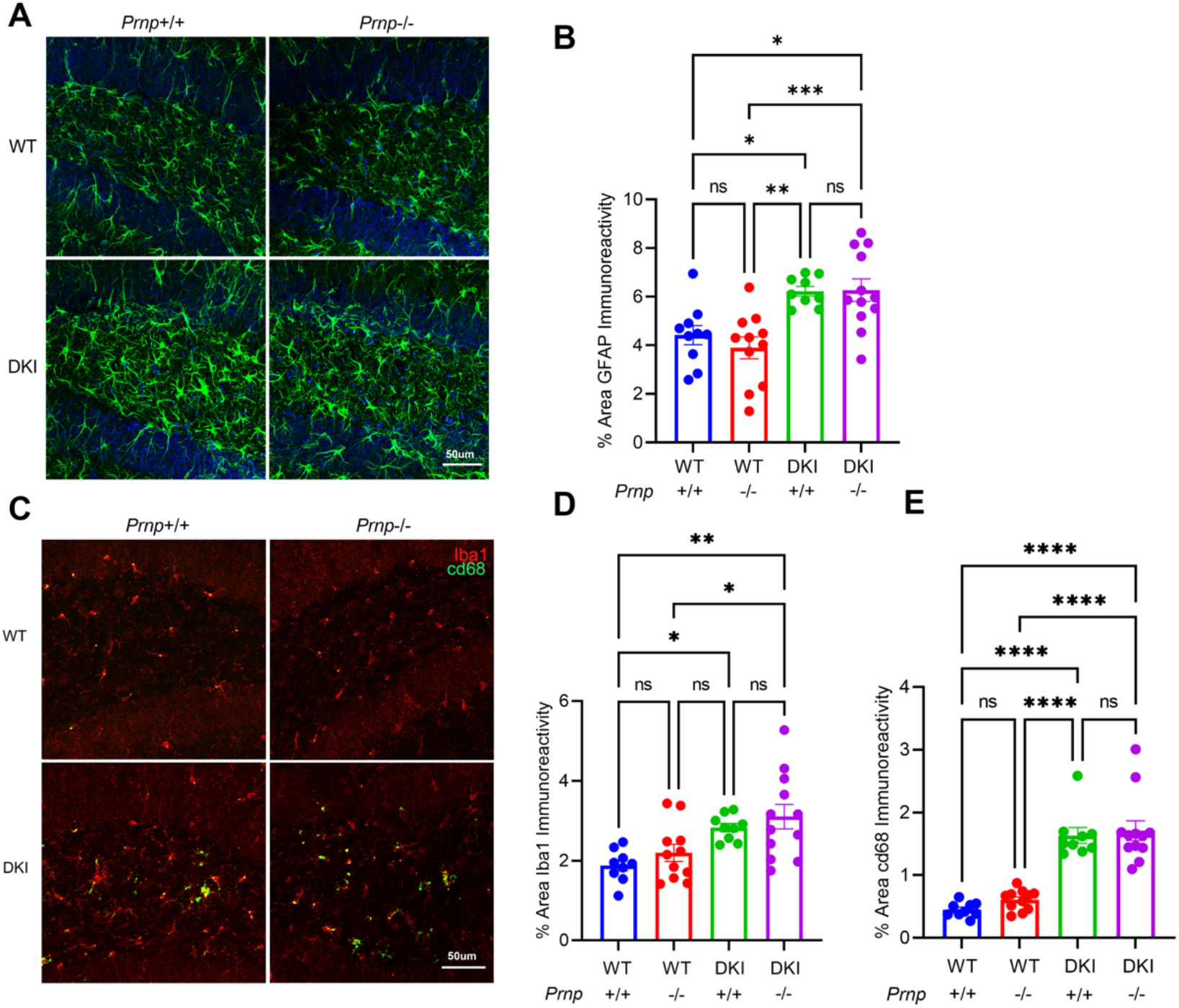
Gliosis in DKI mice is unaffected by *Prnp* deletion. (A) Representative images of GFAP (green) and DAPI (blue) taken in the hippocampus of 10-month-old WT, *Prnp^-/-^*, DKI and DKI; *Prnp^-/-^* mice. Scale bar = 50 μm. (B) Quantification of GFAP immunoreactive area in the hippocampus demonstrates a significant increase in astrocytosis for DKI mice compared to WT, but no effect *Prnp* of knockout. Data are graphed as mean ± SEM, analyzed by ordinary one-way ANOVA with Tukey’s multiple comparisons test *P<0.05,**P<0.01, ***P<0.001, n= 10 for WT, n= 11 for *Prnp^-/-^*, n= 9 for DKI, and n= 12 for DKI; *Prnp^-/-^*. (C) Representative images of Iba1 (red) and CD68 (green) taken in the hippocampus of 10-month-old WT, *Prnp^-/-^*, DKI and DKI; *Prnp^-/-^* mice. Scale bar = 50 μm. (D) Quantification of Iba1 immunoreactive area in the hippocampus demonstrates a significant increase in microgliosis for DKI mice compared to WT, but no effect *Prnp* of knockout. Data are graphed as mean ± SEM, analyzed by ordinary one-way ANOVA with Tukey’s multiple comparisons test *P<0.05,**P<0.01, n= 9 for WT, n= 11 for *Prnp^-/-^*, n= 9 for DKI, and n= 12 for DKI; *Prnp^-/-^*. (E) Quantification of cd68 immunoreactive area in the hippocampus demonstrates a significant increase in activated microglia for DKI mice compared to WT, but no effect *Prnp* of knockout. Data are graphed as mean ± SEM, analyzed by ordinary one-way ANOVA with Tukey’s multiple comparisons test ****P<0.0001, n= 9 for WT, n= 11 for *Prnp^-/-^*, n= 9 for DKI, and n= 12 for DKI; *Prnp^-/-^*.

**Supplemental Figure S4.**
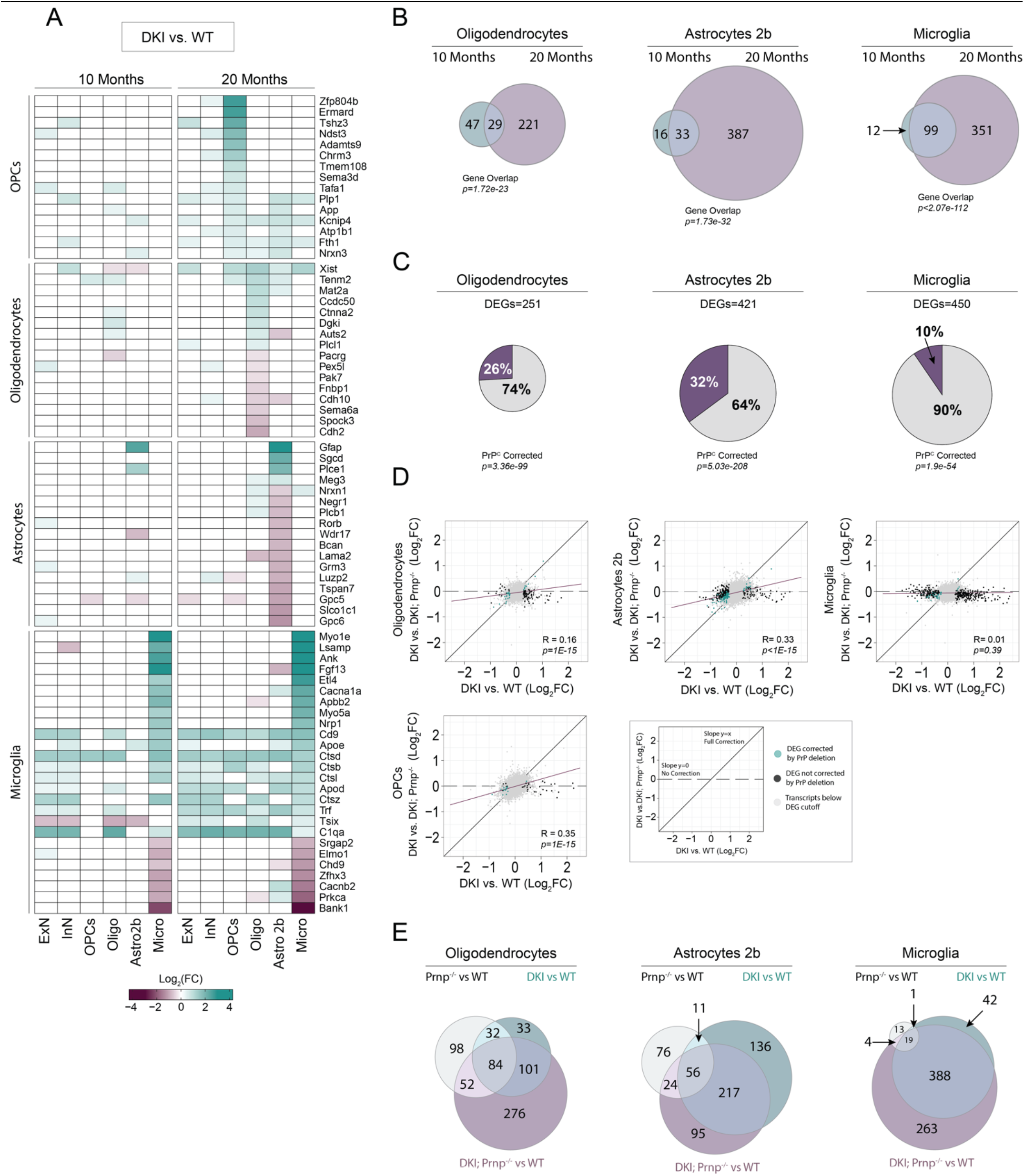
Glial cell activation in a DKI mouse model of AD. **A-E,** AD-associated transcriptional profile of OPCs, oligodendrocytes, astrocytes, and microglia cell populations comparing DKI versus WT samples. **A,** Heatmap of the cellular expression of top differentially expressed glial gene markers at 10 and 20 months. **B,** Venn Diagram comparing shared AD-associated DEGs shared by age-group and cell type. **C,** Pie charts illustrating the percentage of AD-associated DEGs in glial cell populations corrected by PrP^C^ deletion. **B-C,** Sizes of charts corresponds to the number of significant DEGs and statistical significance of gene overlap (Fisher’s Exact Test) is the relative to number of DEGs. **D,** Glial cell-type specific, transcriptome-wide comparisons of AD-associated DEGs and PrP^C^-corrected DEGs. Log2FC between DKI and WT (AD-effect) is plotted along the *x-axis*. Log2FC between DKI and DKI; *Prnp^-/-^* (PrP^C^-effect) is plotted along the *y-axis*. Black points represent significant DEGs with an absolute Log2(FC) > 0.25 and *p-value <0.005*. Colored points represent DEGs also corrected by PrP^C^ deletion. Points along the identity line (x=y) represent genes with equivalent differential expression between WT or DKI; Prnp-/- relative to DKI, indicating complete rescue by PrP^C^ deletion. Points along the line “y=0” reflect genes unaffected by PrP^C^ deletion. The regression line (Pearson’s correlations, purple) represents transcriptome-wide effects of PrP^C^ and *p-values* represents the statistical significance of a non-zero linear regression relationship. **E,** Venn diagrams depicting the number of shared and unique DEGs between *Prnp^-/-^*, DKI, and DKI; *Prnp^-/-^* samples in glial cell populations at 20 months.

**Supplemental Figure S5.**
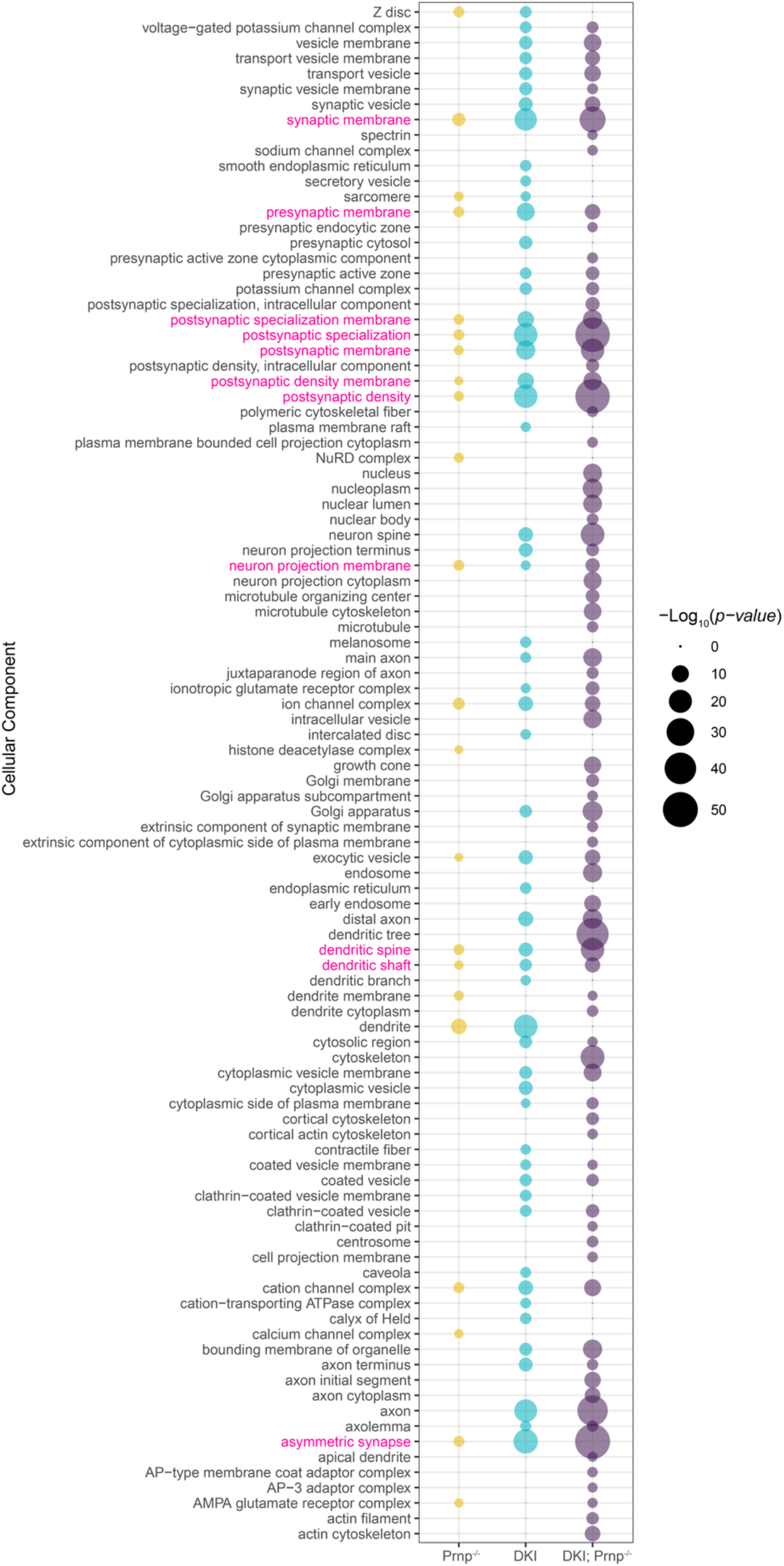
Cellular component analysis of DKI and *Prnp*-dependent DEGs in 10-month neuronal populations. Dot plot of GO Cellular Component terms resulting from the enrichment analysis for DEGs of *Prnp^-/-^*, DKI, and DKI; *Prnp^-/-^* samples as compared to WT. The size of dots indicates the P-value of significance for each term.

**Supplemental Figure S6.**
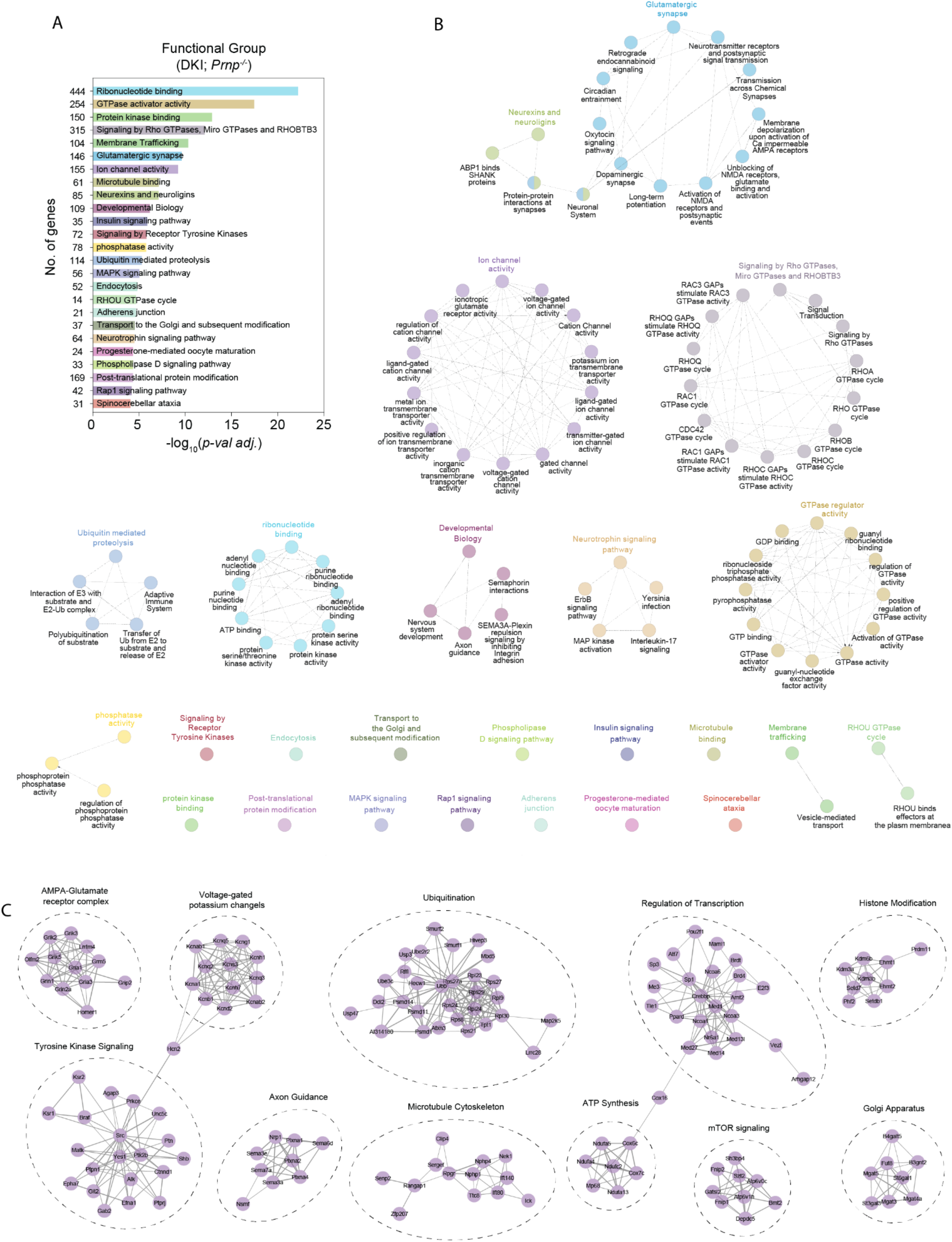
Pathways with altered gene expression dependent on synthetic interaction of DKI model with *Prnp* deletion. **A-C,** Functional pathway enrichment analysis of combined excitatory and inhibitory neuronal DEGs comparing DKI; *Prnp^-/-^* versus WT samples. **B,** Pathway enrichment analysis (GO Molecular Functions, KEGG, and REAC Pathways) with terms (nodes) color coded and organized into functional grouping by gene associations (edges). The most significant leading term is color highlight with its corresponding P-value graphed in **A** along with the total unique gene association count with the respective functional network group. **C,** PPI networks of gene associations grouped by highly enriched functional pathways. The number and thickness of lines represents stronger associations.

## REFERENCES

1. Long JM, Holtzman DM: Alzheimer Disease: An Update on Pathobiology and Treatment Strategies. Cell 2019, 179:312–339.

2. Knopman DS, Amieva H, Petersen RC, Chetelat G, Holtzman DM, Hyman BT, Nixon RA, Jones DT: Alzheimer disease. Nat Rev Dis Primers 2021, 7:33.

3. van Dyck CH, Swanson CJ, Aisen P, Bateman RJ, Chen C, Gee M, Kanekiyo M, Li D, Reyderman L, Cohen S, et al: Lecanemab in Early Alzheimer’s Disease. N Engl J Med 2022.

4. Mintun MA, Lo AC, Duggan Evans C, Wessels AM, Ardayfio PA, Andersen SW, Shcherbinin S, Sparks J, Sims JR, Brys M, et al: Donanemab in Early Alzheimer’s Disease. N Engl J Med 2021, 384:1691–1704.

5. Lauren J, Gimbel DA, Nygaard HB, Gilbert JW, Strittmatter SM: Cellular prion protein mediates impairment of synaptic plasticity by amyloid-beta oligomers. Nature 2009, 457:1128–1132.

6. Smith LM, Strittmatter SM: Binding Sites for Amyloid-beta Oligomers and Synaptic Toxicity. Cold Spring Harbor perspectives in medicine 2017, 7.

7. Smith LM, Kostylev MA, Lee S, Strittmatter SM: Systematic and standardized comparison of reported amyloid-beta receptors for sufficiency, affinity, and Alzheimer’s disease relevance. J Biol Chem 2019, 294:6042–6053.

8. Purro SA, Nicoll AJ, Collinge J: Prion Protein as a Toxic Acceptor of Amyloid-beta Oligomers. Biol Psychiat 2018, 83:358–368.

9. Kostylev MA, Kaufman AC, Nygaard HB, Patel P, Haas LT, Gunther EC, Vortmeyer A, Strittmatter SM: Prion-Protein-interacting Amyloid-beta Oligomers of High Molecular Weight Are Tightly Correlated with Memory Impairment in Multiple Alzheimer Mouse Models. J Biol Chem 2015, 290:17415–17438.

10. Kostylev MA, Tuttle MD, Lee S, Klein LE, Takahashi H, Cox TO, Gunther EC, Zilm KW, Strittmatter SM: Liquid and Hydrogel Phases of PrP(C) Linked to Conformation Shifts and Triggered by Alzheimer’s Amyloid-beta Oligomers. Molecular cell 2018, 72:426–443 e412.

11. Um JW, Nygaard HB, Heiss JK, Kostylev MA, Stagi M, Vortmeyer A, Wisniewski T, Gunther EC, Strittmatter SM: Alzheimer amyloid-beta oligomer bound to postsynaptic prion protein activates Fyn to impair neurons. Nat Neurosci 2012, 15:1227–1235.

12. Gimbel DA, Nygaard HB, Coffey EE, Gunther EC, Lauren J, Gimbel ZA, Strittmatter SM: Memory impairment in transgenic Alzheimer mice requires cellular prion protein. The Journal of neuroscience: the official journal of the Society for Neuroscience 2010, 30:6367–6374.

13. Salazar SV, Gallardo C, Kaufman AC, Herber CS, Haas LT, Robinson S, Manson JC, Lee MK, Strittmatter SM: Conditional Deletion of Prnp Rescues Behavioral and Synaptic Deficits after Disease Onset in Transgenic Alzheimer’s Disease. The Journal of neuroscience: the official journal of the Society for Neuroscience 2017, 37:9207–9221.

14. Cox TO, Gunther EC, Brody AH, Chiasseu MT, Stoner A, Smith LM, Haas LT, Hammersley J, Rees G, Dosanjh B, et al: Anti-PrP(C) antibody rescues cognition and synapses in transgenic alzheimer mice. Ann Clin Transl Neurol 2019, 6:554–574.

15. Gunther EC, Smith LM, Kostylev MA, Cox TO, Kaufman AC, Lee S, Folta-Stogniew E, Maynard GD, Um JW, Stagi M, et al: Rescue of Transgenic Alzheimer’s Pathophysiology by Polymeric Cellular Prion Protein Antagonists. Cell Rep 2019, 26:145–158 e148.

16. Chung E, Ji Y, Sun Y, Kascsak RJ, Kascsak RB, Mehta PD, Strittmatter SM, Wisniewski T: Anti-PrPC monoclonal antibody infusion as a novel treatment for cognitive deficits in an Alzheimer’s disease model mouse. BMC Neurosci 2010, 11:130.

17. Klyubin I, Nicoll AJ, Khalili-Shirazi A, Farmer M, Canning S, Mably A, Linehan J, Brown A, Wakeling M, Brandner S, et al: Peripheral administration of a humanized anti-PrP antibody blocks Alzheimer’s disease Abeta synaptotoxicity. J Neurosci 2014, 34:6140–6145.

18. Haas LT, Salazar SV, Kostylev MA, Um JW, Kaufman AC, Strittmatter SM: Metabotropic glutamate receptor 5 couples cellular prion protein to intracellular signalling in Alzheimer’s disease. Brain 2016, 139:526–546.

19. Spurrier J, Nicholson L, Fang XT, Stoner AJ, Toyonaga T, Holden D, Siegert TR, Laird W, Allnutt MA, Chiasseu M, et al: Reversal of synapse loss in Alzheimer mouse models by targeting mGluR5 to prevent synaptic tagging by C1Q. Science translational medicine 2022, 14:eabi8593.

20. Haas LT, Salazar SV, Smith LM, Zhao HR, Cox TO, Herber CS, Degnan AP, Balakrishnan A, Macor JE, Albright CF, Strittmatter SM: Silent Allosteric Modulation of mGluR5 Maintains Glutamate Signaling while Rescuing Alzheimer’s Mouse Phenotypes. Cell Rep 2017, 20:76–88.

21. Um JW, Kaufman AC, Kostylev M, Heiss JK, Stagi M, Takahashi H, Kerrisk ME, Vortmeyer A, Wisniewski T, Koleske AJ, et al: Metabotropic glutamate receptor 5 is a coreceptor for Alzheimer abeta oligomer bound to cellular prion protein. Neuron 2013, 79:887–902.

22. Brody AH, Strittmatter SM: Synaptotoxic Signaling by Amyloid Beta Oligomers in Alzheimer’s Disease Through Prion Protein and mGluR5. Advances in pharmacology 2018, 82:293–323.

23. Cisse M, Sanchez PE, Kim DH, Ho K, Yu GQ, Mucke L: Ablation of cellular prion protein does not ameliorate abnormal neural network activity or cognitive dysfunction in the J20 line of human amyloid precursor protein transgenic mice. J Neurosci 2011, 31:10427–10431.

24. Saito T, Matsuba Y, Mihira N, Takano J, Nilsson P, Itohara S, Iwata N, Saido TC: Single App knock-in mouse models of Alzheimer’s disease. Nat Neurosci 2014, 17:661–663.

25. Saito T, Mihira N, Matsuba Y, Sasaguri H, Hashimoto S, Narasimhan S, Zhang B, Murayama S, Higuchi M, Lee VMY, et al: Humanization of the entire murine Mapt gene provides a murine model of pathological human tau propagation. J Biol Chem 2019, 294:12754–12765.

26. Manson JC, Clarke AR, Hooper ML, Aitchison L, McConnell I, Hope J: 129/Ola mice carrying a null mutation in PrP that abolishes mRNA production are developmentally normal. Mol Neurobiol 1994, 8:121–127.

27. Sadasivam P, Fang XT, Toyonaga T, Lee S, Xu Y, Zheng MQ, Spurrier J, Huang Y, Strittmatter SM, Carson RE, Cai Z: Quantification of SV2A Binding in Rodent Brain Using [(18)F]SynVesT-1 and PET Imaging. Mol Imaging Biol 2021, 23:372–381.

28. Zhang R, Xue G, Wang S, Zhang L, Shi C, Xie X: Novel object recognition as a facile behavior test for evaluating drug effects in AbetaPP/PS1 Alzheimer’s disease mouse model. J Alzheimers Dis 2012, 31:801–812.

29. Kaufman AC, Salazar SV, Haas LT, Yang J, Kostylev MA, Jeng AT, Robinson SA, Gunther EC, van Dyck CH, Nygaard HB, Strittmatter SM: Fyn inhibition rescues established memory and synapse loss in Alzheimer mice. Ann Neurol 2015, 77:953–971.

30. Bolte S, Cordelieres FP: A guided tour into subcellular colocalization analysis in light microscopy. J Microsc 2006, 224:213–232.

31. Wolf FA, Angerer P, Theis FJ: SCANPY: large-scale single-cell gene expression data analysis. Genome Biol 2018, 19:15.

32. Stuart T, Butler A, Hoffman P, Hafemeister C, Papalexi E, Mauck WM, 3rd, Hao Y, Stoeckius M, Smibert P, Satija R: Comprehensive Integration of Single-Cell Data. Cell 2019, 177:1888–1902 e1821.

33. Shannon P, Markiel A, Ozier O, Baliga NS, Wang JT, Ramage D, Amin N, Schwikowski B, Ideker T: Cytoscape: a software environment for integrated models of biomolecular interaction networks. Genome Res 2003, 13:2498–2504.

34. Bindea G, Mlecnik B, Hackl H, Charoentong P, Tosolini M, Kirilovsky A, Fridman WH, Pages F, Trajanoski Z, Galon J: ClueGO: a Cytoscape plug-in to decipher functionally grouped gene ontology and pathway annotation networks. Bioinformatics 2009, 25:1091–1093.

35. Szklarczyk D, Gable AL, Nastou KC, Lyon D, Kirsch R, Pyysalo S, Doncheva NT, Legeay M, Fang T, Bork P, et al: The STRING database in 2021: customizable protein-protein networks, and functional characterization of user-uploaded gene/measurement sets. Nucleic acids research 2021, 49:D605–D612.

36. Scheff SW, Price DA, Schmitt FA, Mufson EJ: Hippocampal synaptic loss in early Alzheimer’s disease and mild cognitive impairment. Neurobiol Aging 2006, 27:1372–1384.

37. DeKosky ST, Scheff SW: Synapse loss in frontal cortex biopsies in Alzheimer’s disease: correlation with cognitive severity. Ann Neurol 1990, 27:457–464.

38. Le Pichon CE, Valley MT, Polymenidou M, Chesler AT, Sagdullaev BT, Aguzzi A, Firestein S: Olfactory behavior and physiology are disrupted in prion protein knockout mice. Nat Neurosci 2009, 12:60–69.

39. Le Pichon CE, Firestein S: Expression and localization of the prion protein PrP(C) in the olfactory system of the mouse. J Comp Neurol 2008, 508:487–499.

40. Parrie LE, Crowell JAE, Telling GC, Bessen RA: The cellular prion protein promotes olfactory sensory neuron survival and axon targeting during adult neurogenesis. Dev Biol 2018, 438:23–32.

41. Liu Y, Yoo MJ, Savonenko A, Stirling W, Price DL, Borchelt DR, Mamounas L, Lyons WE, Blue ME, Lee MK: Amyloid pathology is associated with progressive monoaminergic neurodegeneration in a transgenic mouse model of Alzheimer’s disease. J Neurosci 2008, 28:13805–13814.

42. Marcyniuk B, Mann DM, Yates PO: The topography of cell loss from locus caeruleus in Alzheimer’s disease. J Neurol Sci 1986, 76:335–345.

43. Busche MA, Hyman BT: Synergy between amyloid-beta and tau in Alzheimer’s disease. Nat Neurosci 2020, 23:1183–1193.

44. Gowrishankar S, Wu Y, Ferguson SM: Impaired JIP3-dependent axonal lysosome transport promotes amyloid plaque pathology. The Journal of cell biology 2017, 216:3291–3305.

45. Gowrishankar S, Yuan P, Wu Y, Schrag M, Paradise S, Grutzendler J, De Camilli P, Ferguson SM: Massive accumulation of luminal protease-deficient axonal lysosomes at Alzheimer’s disease amyloid plaques. Proc Natl Acad Sci U S A 2015, 112:E3699–3708.

46. Yuan P, Zhang M, Tong L, Morse TM, McDougal RA, Ding H, Chan D, Cai Y, Grutzendler J: PLD3 affects axonal spheroids and network defects in Alzheimer’s disease. Nature 2022, 612:328–337.

47. Hong S, Beja-Glasser VF, Nfonoyim BM, Frouin A, Li S, Ramakrishnan S, Merry KM, Shi Q, Rosenthal A, Barres BA, et al: Complement and microglia mediate early synapse loss in Alzheimer mouse models. Science 2016, 352:712–716.

48. Shi Q, Chowdhury S, Ma R, Le KX, Hong S, Caldarone BJ, Stevens B, Lemere CA: Complement C3 deficiency protects against neurodegeneration in aged plaque-rich APP/PS1 mice. Science translational medicine 2017, 9.

49. Erlich P, Dumestre-Perard C, Ling WL, Lemaire-Vieille C, Schoehn G, Arlaud GJ, Thielens NM, Gagnon J, Cesbron JY: Complement protein C1q forms a complex with cytotoxic prion protein oligomers. J Biol Chem 2010, 285:19267–19276.

50. Flores-Langarica A, Sebti Y, Mitchell DA, Sim RB, MacPherson GG: Scrapie pathogenesis: the role of complement C1q in scrapie agent uptake by conventional dendritic cells. J Immunol 2009, 182:1305–1313.

51. Haas LT, Salazar SV, Kostylev MA, Um JW, Kaufman AC, Strittmatter SM: Metabotropic glutamate receptor 5 couples cellular prion protein to intracellular signalling in Alzheimer’s disease. Brain 2015.

52. Palmqvist S, Janelidze S, Quiroz YT, Zetterberg H, Lopera F, Stomrud E, Su Y, Chen Y, Serrano GE, Leuzy A, et al: Discriminative Accuracy of Plasma Phospho-tau217 for Alzheimer Disease vs Other Neurodegenerative Disorders. JAMA 2020, 324:772–781.

53. Haas LT, Salazar SV, Smith LM, Zhao HR, Cox TO, Herber CS, Degnan AP, Balakrishnan A, Macor JE, Albright CF, Strittmatter SM: Silent Allosteric Modulation of mGluR5 Maintains Glutamate Signaling while Rescuing Alzheimer’s Mouse Phenotypes. Cell Reports 2017, 20:76–88.

54. Castle AR, Gill AC: Physiological Functions of the Cellular Prion Protein. Front Mol Biosci 2017, 4:19.

55. Alier K, Li Z, Mactavish D, Westaway D, Jhamandas JH: Ionic mechanisms of action of prion protein fragment PrP(106-126) in rat basal forebrain neurons. J Neurosci Res 2010, 88:2217–2227.

56. Alier K, Ma L, Yang J, Westaway D, Jhamandas JH: Abeta inhibition of ionic conductance in mouse basal forebrain neurons is dependent upon the cellular prion protein PrPC. J Neurosci 2011, 31:16292–16297.

57. Matsubara T, Satoh K, Homma T, Nakagaki T, Yamaguchi N, Atarashi R, Sudo Y, Uezono Y, Ishibashi D, Nishida N: Prion protein interacts with the metabotropic glutamate receptor 1 and regulates the organization of Ca(2+) signaling. Biochem Biophys Res Commun 2020, 525:447–454.

58. Bribian A, Fontana X, Llorens F, Gavin R, Reina M, Garcia-Verdugo JM, Torres JM, de Castro F, del Rio JA: Role of the cellular prion protein in oligodendrocyte precursor cell proliferation and differentiation in the developing and adult mouse CNS. PLoS One 2012, 7:e33872.

59. Klyubin I, Nicoll AJ, Khalili-Shirazi A, Farmer M, Canning S, Mably A, Linehan J, Brown A, Wakeling M, Brandner S, et al: Peripheral Administration of a Humanized Anti-PrP Antibody Blocks Alzheimer’s Disease A Synaptotoxicity. Journal of Neuroscience 2014, 34:6140–6145.

60. Chung E, Ji Y, Sun YJ, Kascsak RJ, Kascsak RB, Mehta PD, Strittmatter SM, Wisniewski T: Anti-PrPC monoclonal antibody infusion as a novel treatment for cognitive deficits in an alzheimer’s disease model mouse. Bmc Neuroscience 2010, 11:130.

61. Kostylev MA, Tuttle MD, Lee S, Klein LE, Takahashi H, Cox TO, Gunther EC, Zilm KW, Strittmatter SM: Liquid and Hydrogel Phases of PrPC Linked to Conformation Shifts and Triggered by Alzheimer’s Amyloid-β Oligomers. Molecular cell 2018, 72:426–443.e412.

